# Independent molecular basis of convergent highland adaptation in maize

**DOI:** 10.1101/013607

**Authors:** Shohei Takuno, Peter Ralph, Kelly Swarts, Rob J. Elshire, Jeffrey C. Glaubitz, Edward S. Buckler, Matthew B. Hufford, Jeffrey Ross-Ibarra

**Author notes:** Present address: SOKENDAI (Graduate university for advanced studies), Hayama, Kanagawa 240-0193, Japan (Shohei Takuno). Corresponding author: Department of Plant Sciences, University of California, Davis, California 95616, USA.

## Abstract

Convergent evolution is the independent evolution of similar traits in different species or lineages of the same species; this often is a result of adaptation to similar environments, a process referred to as convergent adaptation. We investigate here the molecular basis of convergent adaptation in maize to highland climates in Mesoamerica and South America using genome-wide SNP data. Taking advantage of archaeological data on the arrival of maize to the highlands, we infer demographic models for both populations, identifying evidence of a strong bottleneck and rapid expansion in South America. We use these models to then identify loci showing an excess of differentiation as a means of identifying putative targets of natural selection, and compare our results to expectations from recently developed theory on convergent adaptation. Consistent with predictions across a wide parameter space, we see limited evidence for convergent evolution at the nucleotide level in spite of strong similarities in overall phenotypes. Instead, we show that selection appears to have predominantly acted on standing genetic variation, and that introgression from wild teosinte populations appears to have played a role in highland adaptation in Mexican maize.

## Introduction

Convergent evolution occurs when multiple species or populations exhibit similar phenotypic adaptations to comparable environmental challenges (Wood *et al.* 2005; Arendt and Reznick 2008; Elmer and Meyer 2011). Evolutionary genetic analysis of a wide range of species has provided evidence for multiple pathways that lead to convergent evolution. One such route occurs when identical mutations arise independently and fix via natural selection in multiple populations. In humans, for example, malaria resistance due to mutations from Glu to Val at the sixth codon of the *β*-globin gene has arisen independently on multiple unique haplotypes (Currat *et al.* 2002; Kwiatkowski 2005). Convergent evolution can also be achieved when different mutations arise within the same locus yet produce similar phenotypic effects. Grain fragrance in rice appears to have evolved along these lines, as populations across East Asia have similar fragrances resulting from at least eight distinct loss-of-function alleles in the *BADH2* gene (Kovach *et al.* 2009). Finally, convergent evolution may arise from natural selection acting on standing genetic variation in an ancestral population. In the three-spined stickleback, natural selection has repeatedly acted to reduce armor plating in independent colonizations of freshwater environments. Adaptation in these populations occurred both from new mutations as well as standing variation at the *Eda* locus in marine populations (Colosimo *et al.* 2005).

Not all convergent phenotypic evolution is the result of convergent evolution at the molecular level, however. Recent studies of adaptation to high elevation in humans, for example, reveal that the genes involved in highland adaptation are largely distinct among Tibetan, Andean and Ethiopian populations (Bigham *et al.* 2010; Scheinfeldt *et al.* 2012; Alkorta-Aranburu *et al.* 2012). While observations of independent origin may be due to a complex genetic architecture or standing genetic variation, introgression from related populations may also play a role. In Tibetan populations, the adaptive allele at the *EPAS1* locus appears to have arisen via introgression from Denisovans, a related hominid group (Huerta-Sánchez *et al.* 2014). Beyond these examples, however, we still know relatively little about how convergent phenotypic evolution is driven by common genetic changes or the relative frequencies of these different routes of convergent evolution.

The adaptation of maize (*Zea mays* ssp. *mays*) to high elevation environments provides an excellent opportunity to investigate the molecular basis of convergent evolution. Maize was domesticated from the wild teosinte *Zea mays* ssp. *parviglumis* (hereafter *parviglumis*) in the lowlands of southwest Mexico ∼ 9,000 years before present (BP) (Matsuoka *et al.* 2002; Piperno *et al.* 2009; van Heerwaarden *et al.* 2011). After domestication, maize spread rapidly across the Americas, reaching the lowlands of South America and the high elevations of the Mexican Central Plateau by ∼ 6, 000 BP (Piperno 2006), and the Andean highlands by ∼ 4, 000 BP (Perry *et al.* 2006; Grobman *et al.* 2012). The transition from lowland to highland habitats spanned similar environmental gradients in Mesoamerica and S. America (Figure S1) and presented a host of novel challenges that often accompany highland adaptation, including reduced temperature, increased ultraviolet radiation, and reduced partial pressure of atmospheric gases (Körner 2007).

Common garden experiments in Mexico reveal that highland maize has successfully adapted to high elevation conditions (Mercer *et al.* 2008), and phenotypic comparisons between Mesoamerican and S. American populations are suggestive of convergent evolution. Maize landraces (open-pollinated traditional varieties) from both populations share a number of phenotypes not found in lowland populations, including dense macrohairs and stem pigmentation (Wilkes 1977; Wellhausen *et al.* 1957), differences in tassel branch and ear husk number (Brewbaker 2014), and a changed biochemical response to UV radiation (Casati and Walbot 2005). In spite of these shared phenotypes, genetic analyses of maize landraces from across the Americas indicate that the two highland populations are independently derived from their respective lowland populations (Vigouroux *et al.* 2008; van Heerwaarden *et al.* 2011), suggesting that observed patterns of phenotypic similarity are not simply due to recent shared ancestry.

In addition to convergent evolution between maize landraces, a number of lines of evidence suggest convergent evolution in the related wild teosintes. *Zea mays* ssp. *mexicana* (hereafter *mexicana*) is native to the highlands of central Mexico, where it is thought to have occurred since at least the last glacial maximum (Ross-Ibarra *et al.* 2009; Hufford *et al.* 2012a). Phenotypic differences between *mexicana* and the lowland *parviglumis* mirror those between highland and lowland maize (Lauter *et al.* 2004), and population genetic analyses of the two sub-species reveal evidence of natural selection associated with altitudinal differences (Pyhäjärvi *et al.* 2013; Fang *et al.* 2012). Landraces in the highlands of Mexico are often found in sympatry with *mexicana* and gene flow from *mexicana* likely contributed to maize adaptation to the highlands (Hufford *et al.* 2013). No wild *Zea* occur in S. America, and S. American landraces show no evidence of gene flow from Mexican teosinte (van Heerwaarden *et al.* 2011), further suggesting independent origins for altitude-adapted traits.

Here we use genome-wide SNP data from Mesoamerican and S. American landraces to investigate the evidence for convergent evolution to highland environments at the molecular level. We estimate demographic histories for maize in the highlands of Mesoamerica and S. America, then use these models to identify loci that may have been the target of selection in each population. We find a large number of sites showing evidence of selection, consistent with a complex genetic architecture involving many phenotypes and numerous loci. We see little evidence for shared selection across highland populations at the nucleotide or gene level, a result we show is consistent with expectations from recent theoretical work on convergent adaptation (Ralph and Coop 2014a). Instead, our results support a role for adaptive introgression from teosinte in Mexico and highlight the contribution of standing variation to adaptation in both populations.

## Materials and Methods

### Materials and DNA extraction

We included one individual from each of 94 landrace maize accessions from high and low elevation sites in Mesoamerica and S. America (Table S1). Accessions were provided by the USDA germplasm repository or kindly donated by Major Goodman (North Carolina State University). Sampling locations are shown in Figure 1A. Landraces sampled from elevations < 1, 700 m were considered lowland, while accessions from > 1, 700 m were considered highland. Seeds were germinated on filter paper following fungicide treatment and grown in standard potting mix. Leaf tips were harvested from plants at the five leaf stage. Following storage at 80°C overnight, leaf tips were lyophilized for 48 hours. Tissue was then homogenized with a Mini-Beadbeater-8 (BioSpec Products, Inc., Bartlesville, OK, USA). DNA was extracted using a modified CTAB protocol (Saghai-Maroof *et al.* 1984). The quality of DNA was ensured through inspection on a 2% agarose gel and a NanoDrop spectrophotometer (Thermo Scientific, NanoDrop Products, Wilmington, DE, USA).

**Figure 1.**
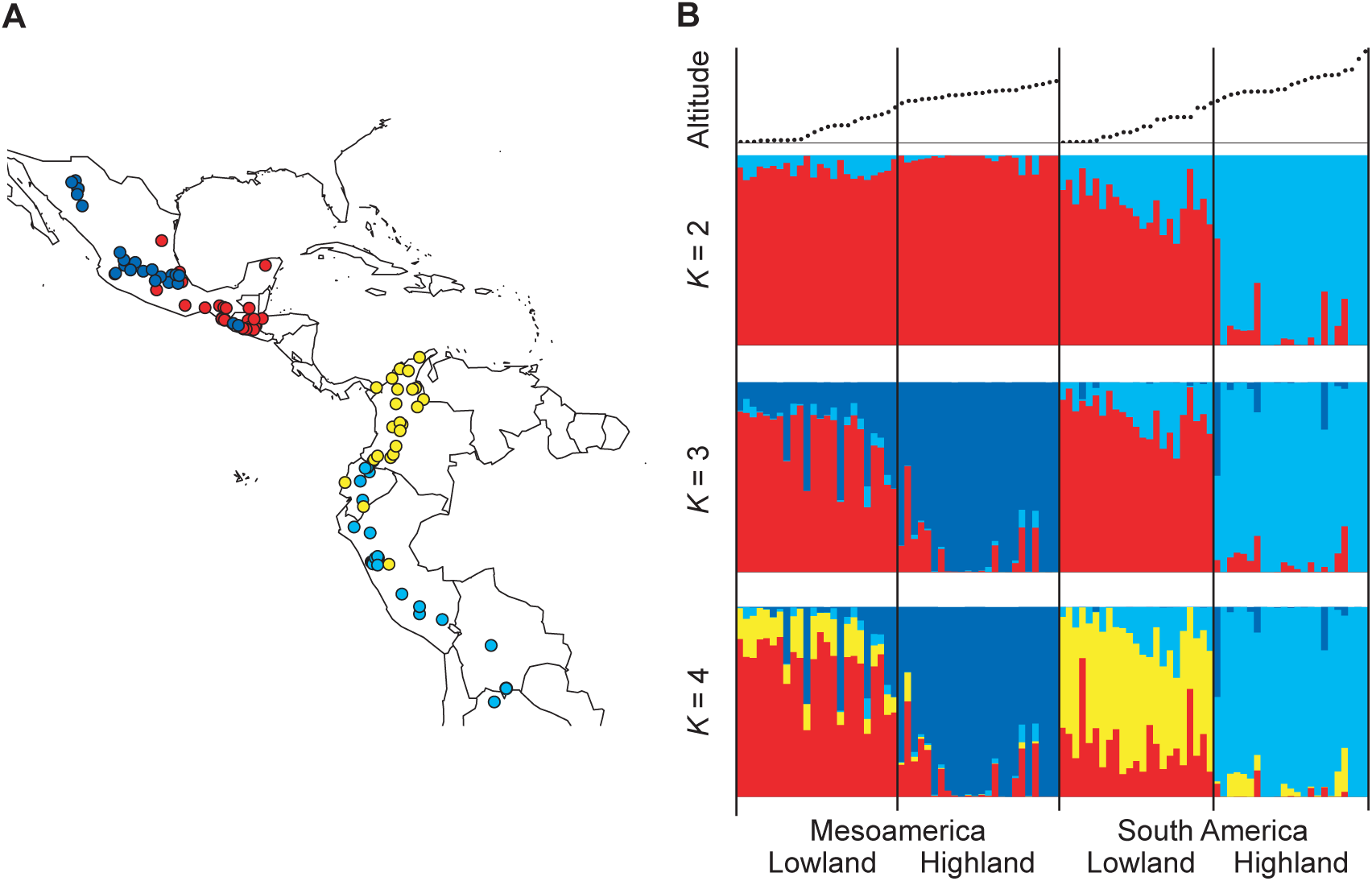
(A) Sampling locations of landraces. Red, blue, yellow and light blue dots represent Mesoamerican lowland, Mesoamerican highland, S. American lowland and S. American highland populations, respectively. (B) Results of STRUCTURE analysis of the maizeSNP50 SNPs with *K* = 2 ∼4. The top panel shows the elevation, ranging from 0 to 4,000 m on the *y*-axes. The colors in *K* = 4 correspond to those in panel (A).

### SNP data

We generated two complementary SNP data sets for the sampled maize landraces. The first set was generated using the Illumina MaizeSNP50 BeadChip platform, including 56,110 SNPs (Ganal *et al.* 2011). SNPs were clustered with the default algorithm of the GenomeStudio Genotyping Module v1.0 (Illumina Inc., San Diego, CA, USA) and then visually inspected and manually adjusted. These data are referred to as “MaizeSNP50” hereafter. This array contains SNPs discovered in multiple ascertainment schemes (Ganal *et al.* 2011), but the vast majority of SNPs come from polymorphisms distinguishing the maize inbred lines B73 and Mo17 (14,810 SNPs) or identified from sequencing 25 diverse maize inbred lines (40,594 SNPs; Gore *et al.* 2009).

The second data set was generated for a subset of 87 of the landrace accessions (Table S1) utilizing high-throughput Illumina sequencing data via genotyping-by-sequencing (GBS; Elshire *et al.* 2011). Genotypes were called using TASSEL-GBS (Glaubitz *et al.* 2014) resulting in 2,848,284 SNPs with an average of 71.3% missing data per individual.

To assess data quality, we compared genotypes at the 7,197 SNPs (229,937 genotypes, excluding missing data) that overlap between the MaizeSNP50 and GBS data sets. While only 0.8% of 173,670 comparisons involving homozygous MaizeSNP50 genotypes differed in the GBS data, 88.6% of 56,267 comparisons with MaizeSNP50 heterozygotes differed, nearly always being reported as a homozygote in GBS. Despite this high heterozygote error rate, the high correlation in allele frequencies between data sets (*r* = 0.89; Figure S2) supports the utility of the GBS data set for estimating allele frequencies.

We annotated SNPs using the filtered gene set from RefGen version 2 of the maize B73 genome sequence (Schnable *et al.* 2009; release 5b.60) from maizesequence.org. We excluded genes annotated as transposable elements (84) and pseudogenes (323) from the filtered gene set, resulting in a total of 38,842 genes.

### Structure analysis

We performed a STRUCTURE analysis (Pritchard *et al.* 2000; Falush *et al.* 2003) using only synonymous and noncoding SNPs from the MaizeSNP50 data due to its low error in identifying heterozygous genotypes. We randomly pruned SNPs closer than 10 kb and assumed free recombination between the remaining SNPs. Alternative distances were tried with nearly identical results. We excluded SNPs in which the number of heterozygous individuals exceeded homozygotes and where the *P*-value for departure from Hardy-Weinberg Equilibrium (HWE) using all individuals was smaller than 0.05 based on a *G*-test. Following these data thinning measures, 17,013 biallelic SNPs remained. We conducted three replicate runs of STRUCTURE using the correlated allele frequency model with admixture for *K* = 2 through *K* = 6 populations, a burn-in length of 50,000 iterations and a run length of 100,000 iterations. Results across replicates were nearly identical.

### Historical population size

We tested three models in which maize was differentiated into highland and lowland populations subsequent to domestication (Figure 2).

**Figure 2.**
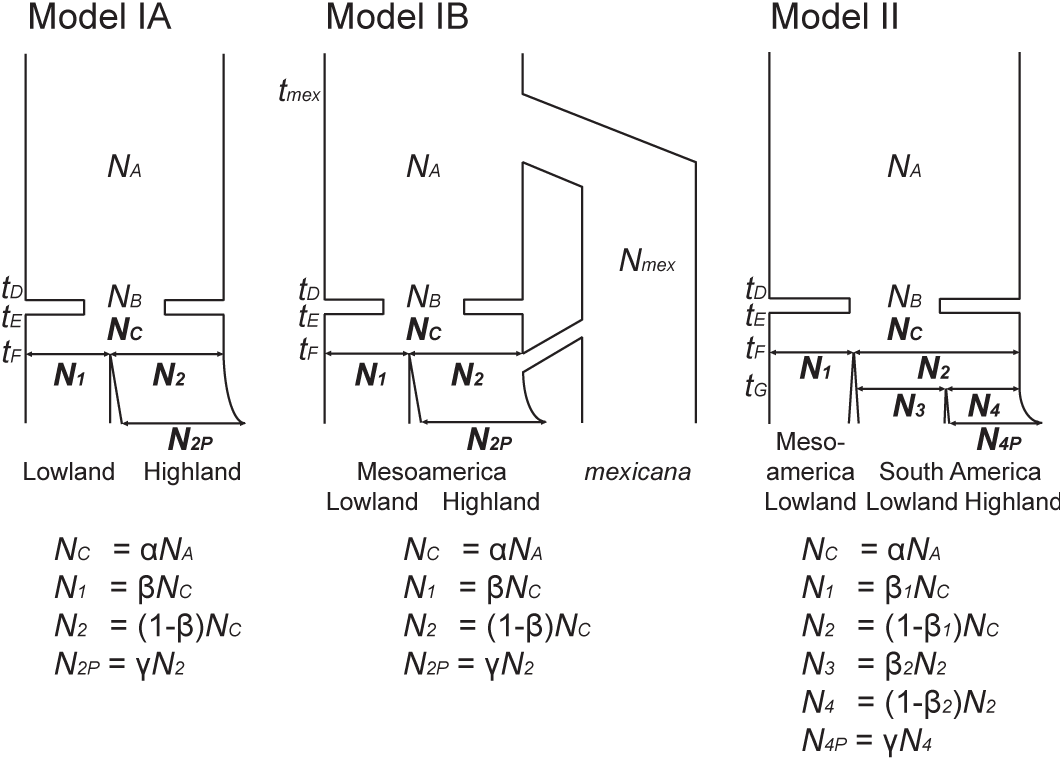
Models of historical population size for lowland and highland populations. Parameters in bold were estimated in this study. See text for details.

We calculated the observed joint frequency distributions (JFDs) using only the GBS data set due to its lower level of ascertainment bias. A subset of synonymous and noncoding SNPs were utilized that had ≥ 15 individuals without missing data in both lowland and highland populations and did not violate HWE. A HWE cut-off of *P* < 0.005 was used for each subpopulation due to our under-calling of heterozygotes.

We obtained similar results under more or less stringent thresholds for significance (*P* < 0.05 ∼ 0.0005; data not shown), though the number of SNPs was very small at *P* < 0.05.

Parameters were inferred with the software *δaδi* (Gutenkunst *et al. 2009),* which uses a diffusion method to calculate an expected JFD and evaluates the likelihood of the data assuming multinomial sampling. We did not use the “full” model that incorporates all four populations because parameter estimation under this model is computationally infeasible.

#### Model IA

This model is applied separately to both the Mesoamerican and the S. American populations. We assume the ancestral diploid population representing *parviglumis* follows a standard Wright-Fisher model with constant size. The size of the ancestral population is denoted by *N*_*A*_. At *t*_*D*_ generations ago, the bottleneck event begins at domestication, and at *t*_*E*_ generations ago, the bottleneck ends. The population size and duration of the bottleneck are denoted by *N*_*B*_ and *t*_*B*_ = *t*_*D*_ − *t*_*E*_, respectively. The population size recovers to *N*_*C*_ = *αN*_*A*_ in the lowlands. Then, the highland population is differentiated from the lowland population at *t*_*F*_ generations ago. The size of the lowland and highland populations at time *t*_*F*_ is determined by a parameter *β* such that the population is divided by *βN*_*C*_ and (1 −*β*)*N*_*C*_; our conclusions hold if we force lowland population size to remain at *N*_*C*_ (data not shown).

We assume that the population size in the lowlands is constant but that the highland population experiences exponential expansion after divergence: its current population size is *γ* times larger than that at *t*_*F.*_

#### Model IB

We expand Model IA for the Mesoamerican populations by incorporating admixture from the teosinte *mexicana* to the highland Mesoamerican maize population. The time of differentiation between *parviglumis* and *mexicana* occurs at *t*_*mex*_ generations ago. The *mexicana* population size is assumed to be constant at *N*_*mex*_. At *t*_*F*_ generations ago, the Mesoamerican highland population is derived from admixture between the Mesoamerican lowland population and a portion *P*_*mex*_ from the teosinte *mexicana*.

#### Model II

The final model includes the Mesoamerican lowland, S. American lowland and highland populations. This model was used for simulating SNPs with ascertainment bias (see below). At time *t*_*F*_, the Mesoamerican and S. American lowland populations are differentiated, and the sizes of populations after splitting are determined by *β*_1_. At time *t*_*G*_, the S. American lowland and highland populations are differentiated, and the sizes of populations at this time are determined by *β*_2_. As in Model IA, the S. American highland population is assumed to experience population growth with the parameter *γ*.

Estimates of a number of our model parameters were available from previous work. *N*_*A*_ was set to 150,000 using estimates of the composite parameter 4*N*_*A*_*µ* ∼ 0.018 from *parviglumis* (Eyre-Walker *et al.* 1998; Tenaillon *et al.* 2001, 2004; Wright *et al*. 2005; Ross-Ibarra *et al*. 2009) and an estimate of the mutation rate *µ* ∼ 3 × 10^−8^ (Clark *et al.* 2005) per site per generation. The severity of the domestication bottleneck is represented by *k* = *N*_*B*_*/t*_*B*_ (Eyre-Walker *et al.* 1998; *Wright et al. 2005), and following Wright et al. (2005) we assumed k* = 2.45 and *t*_*B*_ = 1, 000 generations. Taking into account archaeological evidence (Piperno *et al.* 2009), we assume *t*_*D*_ = 9, 000 and *t*_*E*_ = 8, 000. We further assumed *t*_*F*_ = 6, 000 for Mesoamerican populations in Models IA and IB (Piperno 2006), *t*_*F*_ = 4, 000 for S. American populations in Model IA (Perry *et al.* 2006; Grobman *et al.* 2012), and *t*_*mex*_ = 60, 000, *N*_*mex*_ = 160, 000 (Ross-Ibarra *et al.* 2009), and *P*_*mex*_ = 0.2 (van Heerwaarden *et al.* 2011) for Model IB. For both Models IA and IB, we inferred three parameters (*α*, *β* and *γ*), and, for Model II, we fixed *t*_*F*_ = 6, 000 and *t*_*G*_ = 4, 000 (Piperno 2006; Perry *et al.* 2006; Grobman *et al.* 2012) and estimated the remaining four parameters (*α*, *β*_1_, *β*_2_ and *γ*).

### Population differentiation

We used our inferred models of population size change to generate a null distribution of *F*_*ST*_ from the expected JFD estimated in *δaδi (Gutenkunst et al.* 2009). *The P*-value of a SNP was calculated by *P* (*F*_*ST*_*E*_ ≥ *F*_*ST*_*O*_|*p* ± 0.025) = *P* (*F*_*ST*_*E*_ ≥ *F*_*ST*_*O*_ ∩ *p* ± 0.025)*/P* (*p*± 0.025), where *F*_*ST*_*O*_ and *F*_*ST*_*E*_ are observed and expected *F*_*ST*_ values and *p* ± 0.025 is the set of loci with mean allele frequency across both highland and lowland populations within 0.025 of the SNP in question.

Generating the null distribution of differentiation for the MaizeSNP50 data requires accounting for ascertainment bias. Evaluation of genetic clustering in our data (not shown) coincides with previous work (Hufford *et al.* 2012b) in suggesting that the two inbred lines most important in the ascertainment panel (B73 and Mo17) are most closely related to Mesoamerican lowland maize. We thus added two additional individuals to the Mesoamerican lowland population and generated our null distribution using only SNPs for which the two individuals had different alleles. For model IA in S. America we added two individuals at time *t*_*F*_ to the ancestral population of the S. American lowland and highland populations because the Mesoamerican lowland population was not incorporated into this model. For each combination of sample sizes in lowland and highland populations, we generated a JFD from 10^7^ SNPs using the software ms (Hudson 2002). Then, we calculated *P*-values from the JFD in the same way. We calculated *F*_*ST*_ values for all SNPs that had ≥ 10 individuals with no missing data in all four populations and showed no departure from HWE at the 0.5% (GBS) or 5% (MaizeSNP50) level.

### Haplotype sharing test

We performed a <u>p</u>airwise <u>h</u>aplotype sharing (PHS) test to detect further evidence of selection, following Toomajian *et al.* (2006). To conduct this test, we first imputed and phased the combined SNP data (both GBS and MaizeSNP50) using the fast PHASE software version 1.4.0 (Scheet and Stephens 2006). As a reference for phasing, we used data (excluding heterozygous SNPs) from an Americas-wide sample of 23 partially inbred landraces from the Hapmap v2 data set (Chia *et al.* 2012). We ran fast PHASE with default parameter settings. PHS was calculated for an allele *A* at position *x* by

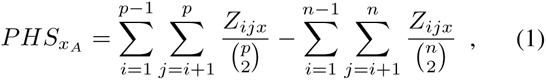

where *n* is the sample size of haploids, *p* is the number of haploids carrying the allele *A* at position *x*, and

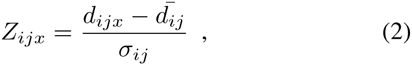

where *d*_*ijx*_ is the genetic distance over which individuals *i* and *j* are identical surrounding position *x*, 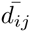 is the genome-wide mean of distances over which individuals *i* and *j* are identical, and *σ*_*ij*_ is the standard deviation of the distribution of distances. Genetic distances were obtained for the MaizeSNP50 data (Ganal *et al.* 2011) and fit using a tenth degree polynomial curve to all SNPs (data not shown).

### Polarizing adaptive alleles

To polarize the ancestral state of alleles and help identify adaptive alleles, we retrieved SNP data from 14 *parviglumis* inbred lines included in the Hapmap v2 data set, using only SNPs with *n* ≥ 10 (Chia *et al.* 2012; Hufford *et al.* 2012b). Alleles were called ancestral if they were at higher frequency in *parviglumis* or uncalled in *parviglumis* but at higher frequency in all populations but one.

For SNPs identified as putative outliers by our *F*_*ST*_ approach, we then used patterns of allele frequency across populations to infer which allele was likely adaptive. For SNPs with a significant *F*_*ST*_ only in Mesoamerica, for example, we characterized them as adaptive if they were at high frequency in one Mesoamerican population (lowland or highland) and low frequency in the other as well as low frequency in *parviglumis* and at most intermediate frequency (or low frequency if missing in *parviglumis*) in S. American populations. SNPs were inferred to show convergent adaptation if they were at high frequency in both highland (or lowland) populations, and at low frequency in the other two populations and *parviglumis*.

### Theoretical evaluation of convergent evolution

We next asked whether the abundance and degree of coincidence of presumably adaptive high-*F*_*ST*_ alleles seen in the SNP data is consistent with what is known about the population history of maize. There are three ways that adaptive alleles could be shared between highland populations: (a) by appearing in both locations as independent, *de novo* mutations; (b) by moving from one highland population to the other by migration; and (c) through convergent selective forces acting on shared standing variation. Here, we provide rough estimates of these rates, and develop in the Appendix more detailed, complementary models that build on the work in Ralph and Coop (2014a) and Ralph and Coop (2014b).

We chose to implement a fairly detailed demographic model. This is because much of the population genetics theory we use relies on universality results that reduce demographic models to relies on universality results that reduce demographic models to two parameters: the dispersal distance (mean parent-offspring distance), and the variance in offspring number. However, these universality results do not hold if either distribution (dispersal or offspring) is sufficiently long-tailed; the detailed model allows us to both get a good idea of what part of parameter space we should focus on, and to verify that the approximation results we use are robust.

To assess the likely importance of (a) and (b), we first evaluate the rate at which we expect an allele that provides a selective advantage at higher elevation to arise by new mutation in or near a highland region (*λ*_mut_), and then use coalescent theory to show that even a highland-adapted allele that was neutral in the lowlands is unlikely to have had time to spread between highland populations under neutral gene flow. It may be more likely that alleles adapted in the highlands are slightly deleterious at lower elevation, consistent with empirical findings in reciprocal transplant experiments in Mexico (Mercer *et al.* 2008); in the Appendix we find the rate at which such an allele already present in the Mesoamerican highlands would transit the intervening lowlands and fix in the Andean highlands. The resulting values depend most strongly on the population density, the selection coefficient, and the rate at which seed is transported long distances and replanted. While long-distance dispersal is certainly possible, evidence from traditional seed systems in Mexico suggests even today it is rare: when farmers exchange seed (a minority of the time) ~ 90% of seed lots come from < 10km away and from a site with altitudinal difference of *<* 50m, although farmers in highland locales exchange seeds over a greater range than average (Bellon *et al.* 2011). We checked the results by evaluating several choices of these parameters as well as with simulations, described in the Appendix. Here we describe the mathematical details; readers may skip to the results without loss of continuity.

#### Demographic model

Throughout, we followed van Heer-waarden *et al.* (2010) in constructing a detailed demographic model for domesticated maize. We assume fields of *N* = 10^5^ plants are replanted each year from *N*_*f*_ = 561 ears, either from completely new stock (with probability *p*_*e*_ = 0.068), from partially new stock (a proportion *r*_*m*_ = 0.2 with probability *p*_*m*_ = 0.02), or otherwise entirely from the same field. Each plant is seed parent to all kernels of its own ears, but can be pollen parent to kernels in many other ears; a proportion *m*_*g*_ = 0.0083 of the pollen-parent kernels are in other fields. Wild-type plants have an average of *µ*_*E*_ = 3 ears per plant, and ears have an average of *N/N*_*f*_ kernels; each of these numbers are Poisson distributed. The mean number of pollen-parent kernels, and the mean number of kernels per ear, is assumed to be (1 + *s*_*b*_) times larger for individuals heterozygous for the selected allele (the fitness of homozygotes is assumed to not affect the probability of establishment). Migration is mediated by seed exchange – when fields are replanted from new stock, the seed is chosen from a random distance away with mean *σ*_*s*_ = 50km, but plants only pollinate other plants belonging to the same village (distance 0). The mean numbers of each category of offspring (seed/pollen; migrant/nonmigrant) are determined by the condition that the population is stable (i.e., wild-type, diploid individuals have on average 2 offspring) except that heterozygotes have on average (1 + *s*_*b*_) offspring that carry the selected allele. Each ear has a small chance of being chosen for replanting, so the number of ears replanted of a given individual is Poisson, and assuming that pollen is well-mixed, the number of pollen-parent kernels is Poisson as well. Each of these numbers of offspring has a mean that depends on whether the field is replanted with new stock, and whether ears are chosen from this field to replant other fields, so the total number of offspring is a mixture of Poissons. These means, and more details of the computations, are found in the Appendix. At the parameter values given, the dispersal distance (mean distance between parent and offspring) is *σ* = 3.5km, and the haploid variance in number of offspring (*ξ*^2^, the variance in number of inherited copies of a chosen parental allele) is between 20 (for wild-type) and 30 (for *s*_*b*_ = 0.1). (Note that in a panmictic population, the offspring variance is approximately the ratio of census size to effective population size, *ξ*^2^ *≈ N/N*_*e*_.)

#### New mutations

The rate at which new mutations appear and fix in a highland population, which we denote *λ*_mut_, is approximately equal to the total population size of the highlands multiplied by the mutation rate per generation and the chance that a single such mutation successfully fixes (i.e., is not lost to drift). The probability that a single new mutant allele providing benefit *s*_*b*_ to heterozygotes at high elevation will fix locally in the high elevation population is approximately 2*s*_*b*_ divided by the haploid variance in offspring number. This can be shown by expanding the generating function near 1, as in Fisher (1922) and Jagers (1975); see Lambert (2006) for more sophisticated models.

Concretely, the probability that a new mutation destined for fixation will arise in a patch of high-elevation habitat of area *A* in a given generation is a function of the density of maize per unit area *ρ*, the selective benefit *s*_*b*_ it provides, the mutation rate *µ*, and the variance in offspring number *ξ*^2^. In terms of these parameters, the rate of appearance is

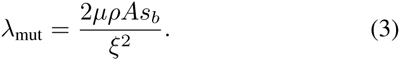

#### Geographic distribution

Throughout we work with populations distributed continuously across geography, with two regions of high elevation, the Mesoamerican and Andean highlands, separated by about 4,000km. The value *A* in equation (3) is the total cultivated area in which the (highland-adapted) alleles in question are beneficial; for estimation of *A* in South America we overlaid raster layers of altitude (www. worldclim.org) and extent of maize cultivation (www. earthstat.org) and calculated the total area of maize cultivated above 1700m using functions in the raster package for R (Hijmans and van Etten 2014).

Of course, the selective benefit of highland alleles is not discrete, but likely changes continuously with altitude, and it may be that the adaptive mutation occurs in a lowland area, subsequently migrating into the highlands. The calculation above does not account for these points, but the approximation is quite good, as verified by exact numerical calculation of the chance of fixation of a mutation as a function of the location where it first appears (see Figure A1); for theoretical treatment see Barton (1987).

**FIGURE A1.**
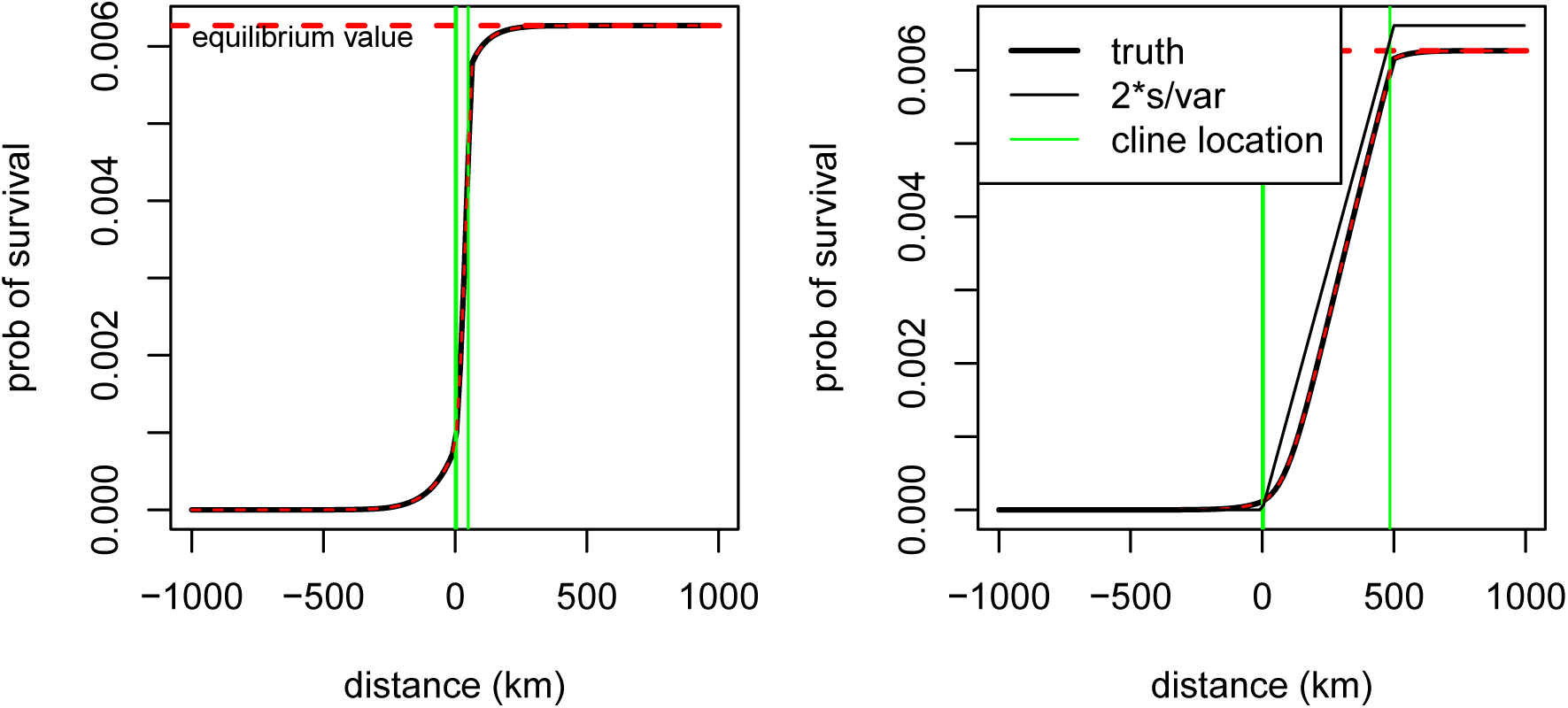
Probability of establishment, as a function of distance along and around an altitudinal cline, whose boundaries are marked by the green lines. **(A)** The parameters above; with cline width 62km; **(B)** the same, except with cline width 500km.

#### Migration

It is harder to intuit a corresponding expression for the chance that an allele established by selection in one highland population moves to the other.

For maize in the Andean highlands to have inherited a highland-adapted allele from the Mesoamerican highlands, those Andean plants must be directly descended from highland Mesoamerican plants that lived more recently than the appearance of the adaptive allele. In other words, the ancestral lineages along which the modern Andean plants have inherited at that locus must trace back to the Mesoamerican highlands. If the allele is neutral in the lowlands, we can treat the movement of these lineages as a neutral process, using the framework of coalescent theory (Wakeley 2005). To do this, we need to follow *all* of the *N ≈* 2.5 × 10^6^ lineages backwards. These quickly coalesce to fewer lineages; but this turns out to not affect the calculation much. Assuming demographic stationarity, the motion of each lineage can be modeled as a random walk, whose displacement after *m* generations has variance *m σ* ^2^, and for large *m* is approximately Gaussian. If we assume that lineages move independently, and *Z*_*n*_ is the distance to the furthest of *n* lineages that 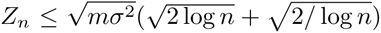 with very high probability (Berman 1964).

Since this depends only on the logarithm of *n*, the number of lineages, the practical upshot of this is that the most distant lineage is very unlikely to be more than about 6 times more distant than the typical lineage, even among 10^7^ lineages. Lineages are not independent, but this only makes this calculation conservative.

## Results

### Samples and data

We sampled 94 maize landraces from four distinct regions in the Americas (Table S1; Figure 1): the lowlands of Mesoamerica (Mexico/Guatemala; *n* = 24) and northern S. America (*n* = 23) and the highlands of Mesoamerica (*n* = 24) and the Andes (*n* = 23). Samples were genotyped using the MaizeSNP50 Beadchip platform (“MaizeSNP50”; *n* = 94) and genotyping-by-sequencing (“GBS”; *n* = 87). After filtering for Hardy-Weinberg genotype frequencies and minimum sample size at least 10 in each of the four populations (see Materials and Methods) 91,779 SNPs remained, including 67,828 and 23,951 SNPs from GBS and MaizeSNP50 respectively.

### Population structure

We performed a STRUCTURE analysis (Pritchard *et al.* 2000; Falush *et al.* 2003) of our landrace samples, varying the number of groups from *K* = 2 to 6 (Figure 1B, Figure S3). Most landraces were assigned to groups consistent with *a priori* population definitions, but admixture between highland and lowland populations was evident at intermediate elevations (∼ 1700m). Consistent with previously described scenarios for maize diffusion (Piperno 2006), we find evidence of shared ancestry between lowland Mesoamerican maize and both Mesoamerican highland and S. American lowland populations. Pairwise *F*_*ST*_ among populations reveals low overall differentiation (Table 1), and the higher *F*_*ST*_ values observed in S. America are consistent with the decreased admixture seen in STRUCTURE. Archaeological evidence supports a more recent colonization of the highlands in S. America (Piperno 2006; Perry *et al.* 2006; Grobman *et al.* 2012), suggesting that the observed differentiation may be the result of a stronger bottleneck during colonization of the S. American highlands.

**Table 1.**
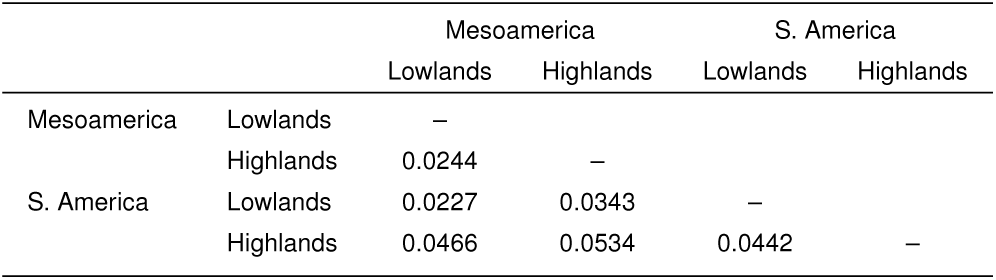
*F*_*ST*_ of synonymous and noncoding GBS SNPs

|  |  | Mesoamerica |  | S. America |  |
| --- | --- | --- | --- | --- | --- |
|  |  | Lowlands | Highlands | Lowlands | Highlands |
| Mesoamerica | Lowlands | – |  |  |  |
|  | Highlands | 0.0244 | – |  |  |
| S. America | Lowlands | 0.0227 | 0.0343 | – |  |
|  | Highlands | 0.0466 | 0.0534 | 0.0442 | – |

### Population differentiation

To provide a null expectation for allele frequency differentiation, we used the joint site frequency distribution (JFD) of low-land and highland populations to estimate parameters of two demographic models using the maximum likelihood method implemented in *δaδi* (Gutenkunst *et al.* 2009). All models incorporate a domestication bottleneck and population differentiation between lowland and highland populations, but differ in their consideration of admixture and ascertainment bias (Figure 2; see Materials and Methods for details). We used published estimates of the strength of the domestication bottleneck (Eyre-Walker et al. 1998; Tenaillon et al. 2004; Wright *et al.* 2005), but confirmed that changing the strength of the bottleneck had little influence on the null distributions of *F*_*ST*_ values (not shown).

Estimated parameter values are listed in Table 2; while the observed and expected JFDs were quite similar for both models, residuals indicated an excess of rare variants in the observed JFDs in all cases (Figure 3). Under both models IA and IB, we found expansion in the highland population in Mesoamerica to be unlikely, but a strong bottleneck followed by population expansion is supported in S. American highland maize in both models IA and II. In Mesoamerica, the likelihood value of model IB was higher than the likelihood of model IA by 850 units of log-likelihood (Table 2), consistent with analyses suggesting a significant role for introgression from *mexicana* during the spread of maize into the highlands (Hufford *et al.* 2013).

**Table 2.** Estimated parameters of population size model

| Mesoamerica | Model IA |  | Model IB |  |
| --- | --- | --- | --- | --- |
|  | Likelihood | –5592.80 | Likelihood | –4654.79 |
| | $N_C$ | 138,000 | $N_C$ | 225,000 |
| | $N_1$ | 52,440 | $N_1$ | 171,000 |
| | $N_2$ | 85,560 | $N_2$ | 54,000 |
| | $N_{2P}$ | 85,560 | $N_{2P}$ | 54,000 |
| S. America | Model IA |  | Model II |  |
|  | Likelihood | –3855.28 | Likelihood | –8044.71 |
| | $N_C$ | 78,000 | $N_C$ | 150,000 |
| | $N_1$ | 75,660 | $N_1$ | 96,000 |
| | $N_2$ | 2,340 | $N_2$ | 54,000 |
| | $N_{2P}$ | 205,920 | $N_3$ | 51,300 |
| | | | $N_4$ | 2,700 |
| | | | $N_{4P}$ | 145,800 |

**Figure 3.**
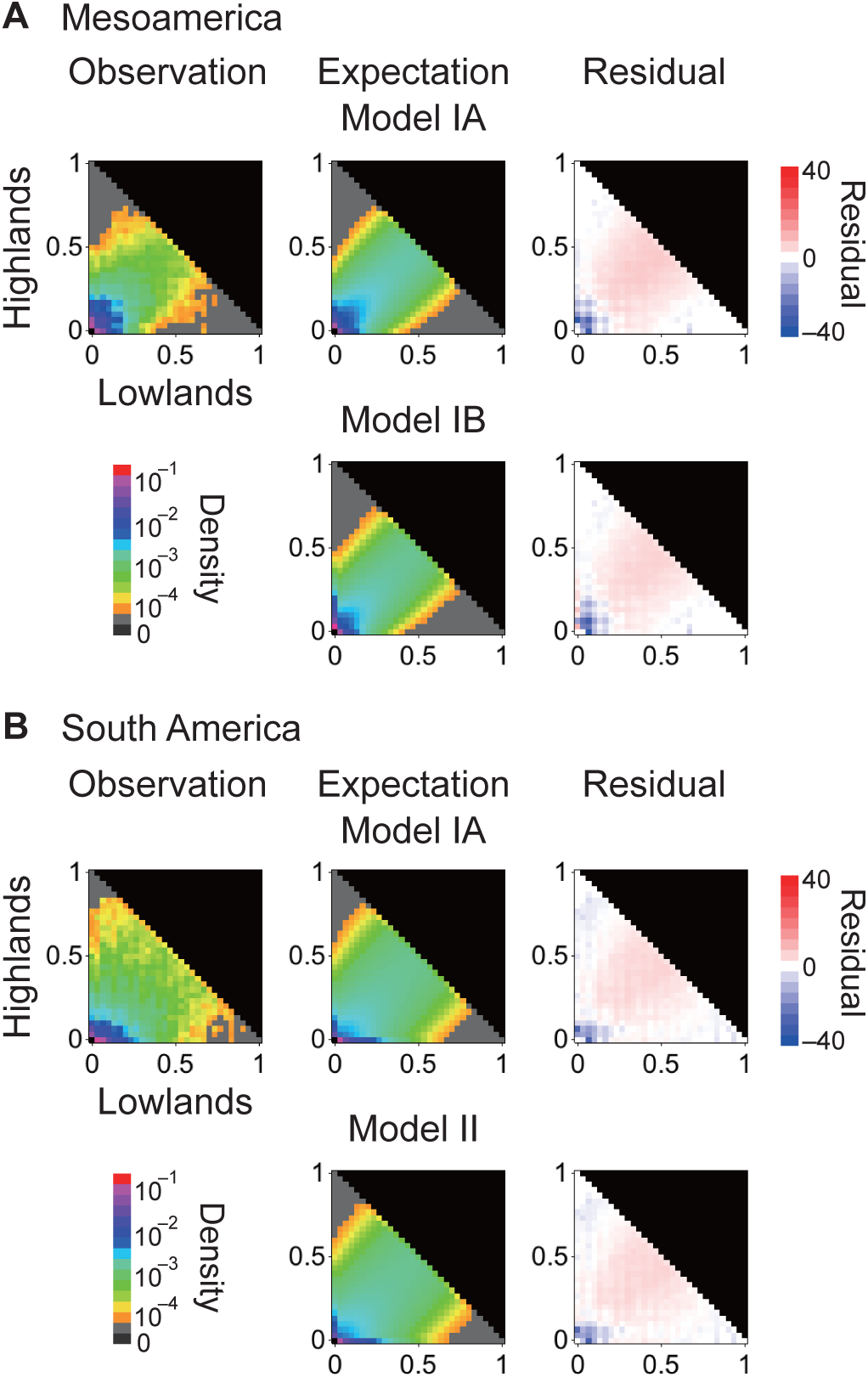
Observed and expected joint distributions of minor allele frequencies in lowland and highland populations in (A) Mesoamerica and (B) S. America. Residuals are calculated as 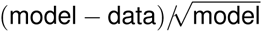.

Comparisons of our empirical *F*_*ST*_ values to the null expectation simulated under our demographic models allowed us to identify significantly differentiated SNPs between lowland and highland populations. In all cases, observed *F*_*ST*_ values were quite similar to those generated under our null models (Figure S4), and model choice had little impact on the distribution of estimated *P*-values (Figure S5). We show results under Model IB for Mesoamerican populations and Model II for S. American populations. We chose *P* < Residual 0.01 as the cut-off for significant differentiation between lowland and highland populations, and identified 687 SNPs in Mesoamerica (687/76,989=0.89%) and 409 SNPs in S. America (409/63,160=0.65%) as significant outliers (Figure 4). All results were qualitatively identical with different cutoff values (0.05 or 0.001; data not shown). SNPs with significant *F*_*ST*_*P*-values were enriched in intergenic regions rather than protein coding regions (60.0% vs. 47.9%, Fisher’s Exact Test *P* < 10^−7^ for Mesoamerica; 62.0% vs. 47.8%, FET *P* < 10^−5^ for S. America).

**Figure 4.**
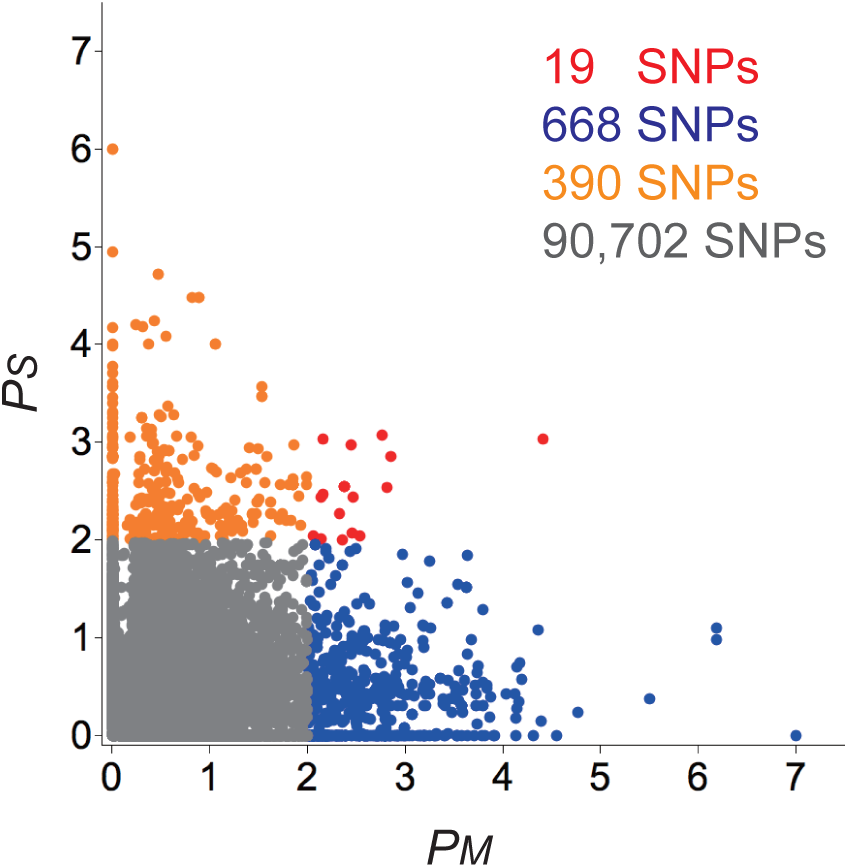
Scatter plot of log10 *P*-values of observed *F*_*ST*_ values based on simulation from estimated demographic models. *P*-values are shown for each SNP in both Mesoamerica (Model IB; *P*_*M*_ on *x*-axis) and S. America (Model II; *P*_*S*_ on *y*-axis). Red, blue, orange and gray dots represents SNPs showing significance in both Mesoamerica and S. America, only in Mesoamerica, only in S. America, or in neither region, respectively. The number of SNPs in each category is shown in the same color as the points.

### Patterns of adaptation

Given the historical spread of maize from an origin in the lowlands, it is tempting to assume that the observation of significant population differentiation at a SNP should be primarily due to an increase in frequency of adaptive alleles in the highlands. To test this hypothesis, we sought to identify the adaptive allele at each locus using comparisons between Mesoamerica and S. America as well as to *parviglumis* (see Methods). Consistent with predictions, we infer that differentiation at 72.3% (264) and 76.7% (230) of SNPs in Mesoamerica and S. America is due to adaptation in the highlands after excluding SNPs with ambiguous patterns likely due to recombination (Table S2).

As further evidence of selection, we asked whether alleles showing excess differentiation also exhibit longer haplotypes than expected. We calculated the empirical quantile of the pairwise haplotype score from Toomajian *et al.* (2006) for each putatively adaptive SNP as the proportion of all SNPs at a similar frequency with PHS scores greater than or equal to the PHS score observed at the focal SNP (Table S2). If *F*_*ST*_ outliers have indeed been targeted by selection in a particular population, we expect this empirical quantile to be smaller (i.e., fewer random SNPs of similar frequency have as large a PHS score) than in other populations. Indeed, we find that SNPs identified as putatively adaptive in each of the four populations show smaller empirical PHS quantiles more often than the 50% expected by chance (Table S2).

Convergent evolution at the nucleotide level should be reflected in an excess of SNPs showing significant differentiation between lowland and highland populations in both Mesoamerica and S. America. Although the 19 SNPs showing *F*_*ST*_ *P*-values *<* 0.01 in both Mesoamerica (*P*_*M*_) and S. America (*P*_*S*_) is statistically greater than the ≈ 5 expected (48, 370 × 0.01 %#x00D7; 0.01 ≈ 4.8; *χ*^2^-test, *P ≪* 0.001), it nonetheless represents a small fraction (≈7 − 8%) of all SNPs showing evidence of selection. This paucity of shared selected SNPs does not appear to be due to our demographic model: a simple outlier approach based using the 1% highest *F*_*ST*_ values finds no shared adaptive SNPs between Mesoamerican and S. American highland populations. For 13 of 19 SNPs showing putative evidence of shared selection we could use data from *parviglumis* to infer whether these SNPs were likely selected in lowland or highland conditions (see Methods). Surprisingly, SNPs identified as shared adaptive variants more frequently showed segregation patterns consistent with lowland (10 SNPs) rather than highland adaptation (2 SNPs).

We also investigated how often different SNPs in the same gene may have been targeted by selection. To search for this pattern, we considered all SNPs within 10kb of a transcript as part of the same gene, excluding SNPs in an miRNA or second transcript. We classified SNPs showing significant *F*_*ST*_ in Mesoamerica, S. America or in both regions into 778 genes. Of these, 485 and 277 genes showed Mesoamerica-specific and SA-specific significant SNPs, while 14 genes contained at least one SNP with a pattern of differentiation suggesting convergent evolution and 2 genes contained both Mesoamerica-specific and SA-specific significant SNPs. Overall, however, fewer genes showed evidence of convergent evolution than expected by chance (permutation test; *P* < 10^−5^).

Finally, we tested whether genes showing evidence of selection in both highland populations were enriched for particular metabolic pathways using data on 481 metabolic pathways from the MaizeCyc database (ver. 2.2; We found 92 pathways that include a selected gene from only one of the highland populations, but no significant excess of shared pathways: only 32 pathways included a selected gene in both populations (*P* = 0.0961; Table S3). Despite similar phenotypes and environments, we thus see little evidence for convergent evolution at the SNP, gene, and metabolic-pathway levels.

### Comparison to theory

Given the limited empirical evidence for convergent evolution at the molecular level, we took advantage of recent theoretical efforts (Ralph and Coop 2014a) to assess the degree of convergence expected under a spatially explicit population genetic model (see Materials and Methods). Using current estimates of maize cultivation in S. America, we find a 270, 200km^2^ area in which maize is cultivated in ≥ 1% of the land area, for a total area of cultivation of ≈ 600, 000ha. At a planting density of *ρ ≈* 20, 000 plants per hectare, this gives a total maize population of ≈12 billion. Assuming an offspring variance of *ξ*^2^ = 30, we can then compute the waiting time *T*_mut_ = 1*/λ*_mut_ for a new beneficial mutation to appear and fix. If we assume an average selection coefficient of *s*_*b*_ = 10^−5^ for each mutation, a single-base mutation with mutation rate *µ* = 3 × 10^−8^ (Clark *et al.* 2005) would take an expected 4,162 generations to appear and fix. Our estimate of the maize population size uses the land area currently under cultivation and is likely an overestimate; *T*_mut_ scales linearly with the population size and lower estimates of *A* will thus increase *T*_mut_ proportionally. However, because *T*_mut_ also scales approximately linearly with both the selection coefficient and the mutation rate, strong selection and the existence of multiple equivalent mutable sites could reduce this time. For example, if any one of 10 sites within a gene were to have an equivalent selective benefit of *s*_*b*_ = 10^−4^, *T*_mut_would be reduced to 42 generations assuming constant *A* over time.

Gene flow between highland regions could also generate patterns of shared adaptive SNPs. The coalescent calculations described above suggest that highland area today is unlikely to draw any ancestry from a region more than 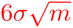 kilometers away from *m* generations ago in any part of the genome that is neutral in the lowlands. Our estimated dispersal of *σ* = 3.5km thus provides an estimate of 1,328km. The Mesoamerican and Andean highlands are approximately 4,000 km apart, and neutral alleles are therefore not expected to transit between the Mesoamerican and Andean highlands within 4,000 generations. Changing the typical distance over which farmers share seed by a factor of 10 would change this conclusion, but data from field surveys do not lend support to such high dispersal distances (Bellon *et al.* 2011).

These results for neutral alleles put a lower bound on the time for deleterious alleles to transit as well, suggesting that we should not expect even weakly deleterious alleles (e.g., *s*_*m*_ = 10^−5^) to have moved between highlands. We expect many of the alleles adaptive in the highlands to be deleterious in the lowlands, and analyze this case in more detail in the Appendix.

Taken together, these theoretical considerations suggest that any alleles beneficial in the highlands that are neutral or deleterious in the lowlands and shared by both the Mesoamerican and S. American highlands would have been present as standing variation in both populations, rather than passed between them.

### Alternative routes of adaptation

The lack of both empirical and theoretical support for convergent adaptation at SNPs or genes led us to investigate alternative patterns of adaptation.

We first sought to understand whether SNPs showing high differentiation between the lowlands and the highlands arose primarily via new mutations or were selected from standing genetic variation. We found that putatively adaptive variants identified in both Mesoamerica and S. America tended to segregate in both the lowland population (85.3% vs. 74.8% in Mesoamerica (Fisher’s exact test *P* < 10^−9^ and 94.8% vs 87.4% in S. America, *P* < 10^−4^) and *parviglumis* (78.3% vs. 72.2% in Mesoamerica (Fisher’s exact test *P* <0.01 and 80.2% vs 72.8% in S. America, *P* < 0.01) more often than other SNPs of similar mean allele frequency.

While maize in highland Mesoamerica grows in sympatry with the highland teosinte *mexicana*, maize in S. America is outside the range of wild *Zea* species, leading to a marked difference in the potential for adaptive introgression from wild relatives. Pyhäjärvi *et al.* (2013) recently investigated local adaptation in *parviglumis* and *mexicana* populations, characterizing differentiation between these subspecies using an outlier approach. Genome-wide, only a small proportion (2–7%) of our putatively adaptive SNPs were identified by Pyhäjärvi *et al.* (2013), though these numbers are still in excess of expectations (Fisher’s exact test *P* < 10^−3^ for S. America and *P* < 10^−8^ for Mesoamerica; Table S4). The proportion of putatively adaptive SNPs shared with teosinte was twice as high in Mesoamerica, however, leading us to evaluate the contribution of introgression from *mexicana* (Hufford *et al.* 2013) in patterning differences between S. American and Mesoamerican highlands.

The proportion of putatively adaptive SNPs in introgressed regions of the genome in highland maize in Mesoamerica was nearly four times higher than found in S. America (FET P < 10^−11^), while differences outside introgressed regions were much smaller (7.5% vs. 6.2%; Table S5). Furthermore, of the 77 regions identified as introgressed in Hufford *et al.* (2013), more than twice as many contain at least one *F*_*ST*_ outlier in Mesoamerica as in S. America (23 compared to 9, one-tailed Z-test *P* = 0.0027). Excluding putatively adaptive SNPs, mean *F*_*ST*_ between Mesoamerica and S. America is only slightly higher in introgressed regions (0.032) than across the rest of the genome (0.020), suggesting the enrichment of high *F*_*ST*_ SNPs seen in Mesoamerica is not simply due to neutral introgression of a divergent teosinte haplotype.

## Discussion

Our analysis of diversity and population structure in maize landraces from Mesoamerica and S. America points to an independent origin of S. American highland maize, in line with earlier archaeological (Piperno 2006;Perry *et al.* 2006;Grobman *et al.* 2012) and genetic (van Heerwaarden *et al.*2011) work. We use our genetic data to fit a model of historical population size change, and find evidence of a strong bottleneck followed by expansion in the highlands of S. America. We identified SNPs deviating from patterns of allele frequencies determined by our demographic model as loci putatively under selection for highland adaptation.

Though the rapid decay of linkage disequilibrium in maize (Figure S6) makes it likely we have identified only a subset of selected loci (Tiffin and Ross-Ibarra 2014), several lines of evidence suggest our results are likely representative of genome-wide patterns. SNPs identified as *F*_*ST*_ outliers by our method show evidence of longer haplotypes and patterns of among-population allele frequency consistent with adaptation (Table S2). Consistent with previous work suggesting adaptive introgression from teosinte, the Mesoamerican highland population shares a larger proportion of SNPs identified as adaptive in teosinte (Pyhäjärvi *et al.*2013). We also see more *F*_*ST*_ outliers Mesoamerica in regions introgressed from teosinte and which overlap with QTL for differences between *parviglumis* and *mexicana* (Lauter *et al.* 2004;Hufford *et al.*2013). Finally, though our SNP data are enriched in low-copy genic regions, our results are consistent with both GWAS in maize (Wallace et al. 2014) and local adaptation in teosinte (Pyhäjärvi et al. 2013) in finding an excess of putatively adaptive SNPs in intergenic regions of the genome.

Although our data identify hundreds of loci that may have been targeted by natural selection in Mesoamerica and S. America, fewer than 1.8% of SNPs and 2.1% of genes show evidence for convergent evolution between the two highland populations. This relative lack of convergent evolution is concordant with recently developed theory (Ralph and Coop 2014a), which applied to this system suggests that convergent evolution involving identical nucleotide changes is unlikely to have occurred in the time since highland colonization through either recurrent mutation or migration across Central America via seed sharing. These results are generally robust to variation in most of the parameters, but are sensitive to gross misestimation of some of the parameters – for example if seed sharing was common over distances of hundreds of kilometers. The modeling highlights that our outlier approach may not detect traits undergoing convergent evolution if the genetic architecture of the trait is such that mutation at a large number of nucleotides would have equivalent effects on fitness (i.e. adaptive traits have a large mutational target). While QTL analysis suggests that some of the traits suggested to be adaptive in highland conditions may be determined by only a few loci (Lauter et al. 2004), others such as flowering time (Buckler et al. 2009) are likely to be the result of a large number of loci, each with small and perhaps similar effects on phenotype. Future quantitative genetic analysis of highland traits using genome-wide association methods may prove useful in searching for the signal of selection on such highly quantitative traits.

Our observation of little convergent evolution is also consistent with the possibility that much of the adaptation to highland environments made use of standing genetic variation in lowland populations. Indeed, we find that as much as 90% of the putatively adaptive variants in Mesoamerica and S. America are segregating in lowland populations, and the vast majority are also segregating in teosinte. Selection from standing variation should be common when the scaled mutation rate *θ* (product of the effective population size, mutation rate and target size) is greater than 1, as long as the scaled selection coefficient N s (product of the effective population size and selection coefficient) is reasonably large (Hermisson and Pennings 2005). Estimates of *θ* from synonymous nucleotide diversity in maize (Tenaillon et al. 2004; Wright et al. 2005; Ross-Ibarra et al. 2009), suggest that adaptation from standing genetic variation may be likely for target sizes larger than a few hundred nucleotides. In maize, such a scenario has been recently shown for the locus grassy tillers1 (Wills et al. 2013), at which adaptive variants in both an upstream control region and the 3’ UTR are segregating in teosinte but show evidence of recent selection in maize, presumably due to the effects of this locus on branching and ear number.

Both our empirical and theoretical results suggest that adaptation to high elevation probably occurred through some combination of selection on standing variation and independent de novo mutation at highly quantitative traits. Because cultivated maize has retained high levels of diversity, much of the ancestral variation present in the populations that founded each of the two highlands was likely shared, allowing for the possibility of shared signals due to selection on the same ancestral variants. However, initial frequencies of alleles present as standing variation will be highly stochastic, leading to a significant role of chance in which alleles are selected, as well the strength of the signal of *F*_*ST*_. This is particularly true for alleles likely to be adaptive in the highlands and thus weakly deleterious in lowland populations, as these should be rare in individual populations. Epistasis could make it even less likely that the same allele is shared between regions.

Overall, our results highlight the complexity of studying convergent evolution for quantitative traits in highly diverse species. Our future efforts will take advantage of reciprocal transplant experiments to identify specific phenotypes under selection. We are also developing mapping populations in both Mesoamerica and South America that should allow identification of genomic regions underlying phenotypes of interest and estimation of the proportion of adaptive variation shared between populations.

## Acknowledgements

We appreciate the helpful comments of P. Morrell and members of the Ross-Ibarra lab and Coop labs. This project was supported by Agriculture and Food Research Initiative Competitive Grant 2009-01864 from the USDA National Institute of Food and Agriculture as well as funding from National Science Foundation grants IOS-1238014 (to JRI) and DBI-1262645 (to PLR).

## Appendix

### Demographic modeling

Throughout we use in many ways the *branching process approximation* – if an allele is locally rare, then for at least a few generations, the fates of each offspring are nearly independent. So, if the allele is locally deleterious, the total numbers of that allele behave as a subcritical branching process, destined for ultimate extinction. On the other hand, if the allele is advantageous, it will either die out or become locally common, with its fate determined in the first few generations. If the number of offspring of an individual with this allele is the random variable *X*, with mean 𝔼 [*X*] = 1 + *s* (selective advantage *s >* 0), variance Var[*X*] = *ξ*^2^, and ℙ {*X* = 0} *>* 0 (some chance of leaving no offspring), then the probability of local nonextinction *p*_*_ is approximately *p*_*_ *≈* 2*s/ξ*^2^ to a second order in *s*. The precise value can be found by defining the generating function Φ(*u*) = 𝔼 [*u*^*X*^]; the probability of local nonextinction *p*_*_ is the minimal solution to Φ(1 *−u*) = 1 *− u*. (This can be seen because: 1 *− p*_*_ is the probability that an individual’s family dies out; this is equal to the probability that the families of all that individuals’ children die out; since each child’s family behaves independently, if the individual has *x* offspring, this is equal to (1 *− p*_*_)^*x*^; and Φ(1− *p*_*_) = 𝔼 [(1 *− p*_*_)^*X*^].)

If the selective advantage (*s*) depends on geographic location, a similar fact holds: index spatial location by *i ∈* 1*, …, n*, and for *u* = (*u*_1_*, u*_2_*,…, u*_*n*_) define the functions 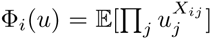where *X*_*ij*_ is the (random) number of offspring that an individual at *i* produces at location *j*. Then *p*_*_= (*p*_*_1*,… p*_*\*n*_), the vector of probabilities that a new mutation at each location eventually fixes, is the minimal solution to Φ(1 *− p_*_*) = 1 *− p_*_*, i.e. Φ_*i*_ (1 *− p*_*\**_) = 1 *− p*_*\*i*_.

Here we consider a linear habitat, so that the selection coefficient at location *ℓ*_*i*_ is *s*_*i*_ = min(*s*_*b*_, max(−*s*_*d*_*, αℓ*_*i*_)). There does not seem to be a nice analytic expression for *p*_*\**_ in this case, but since 1 −*p*_*\**_ is a fixed point of Φ, the solution can be found by iteration: 1 −*p*_*\**_ = lim_*n→∞*_ Φ^*n*^(*u*) for an appropriate starting point *u*.

#### Maize model

The migration and reproduction dynamics we use are taken largely from van Heerwaarden *et al.* (2010). On a large scale, fields of *N* plants are replanted each year from *N*_*f*_ ears, either from completely new stock (with probability *p*_*e*_), from partially new stock (a proportion *r*_*m*_ with probability *p*_*m*_), or entirely from the same field. Plants have an average of *μ*_*E*_ ears per plant, and ears have an average of *N/N*_*f*_ kernels; so a plant has on average *μ*_*E*_*N/N*_*f*_ kernels, and a field has on average *μ*_*E*_*N* ears and *μ*_*E*_*N* ^2^*/N*_*f*_ kernels. We suppose that a plant with the selected allele is pollen parent to (1 + *s*)*μ*_*E*_*N/N*_*f*_ kernels, and also seed parent to (1 + *s*)*μ*_*E*_*N/N*_*f*_ kernels, still in *μ*_*E*_ ears. The number of offspring a plant has depends on how many of its offspring kernels get replanted. Some proportion *m*_*g*_ of the pollen-parent kernels are in other fields, and may be replanted; but with probability *p*_*e*_ no other kernels (i.e. those in the same field) are replanted. Otherwise, with probability 1*− p*_*m*_ the farmer chooses *N*_*f*_ of the ears from this field to replant (or, (1*−r*_*m*_)*N*_*f*_ of them, with probability *p*_*m*_); this results in a mean number *N*_*f*_ */N* (or, (1*−r*_*m*_)*N*_*f*_ */N*) of the plant’s ears of seed children being chosen, and a mean number 1 + *s* of the plant’s pollen children kernels being chosen. Furthermore, the field is used to completely (or partially) replant another’s field with chance *p*_*e*_*/*(1 − *p*_*e*_) (or *p*_*m*_); resulting in another *N*_*f*_ */N* (or *r*_*m*_*N*_*f*_ */N*) ears and 1 + *s* (or *r*_*m*_(1 + *s*)) pollen children being replanted elsewhere. Here we have assumed that pollen is well-mixed within a field, and that the selected allele is locally rare. Finally, we must divide all these offspring numbers by 2, since we look at the offspring carrying a particular haplotype, not of the diploid plant’s genome.

The above gives mean values; to get a probability model we assume that every count is Poisson. In other words, we suppose that the number of pollen children is Poisson with random mean *λ*_*P*_, and the number of seed children is a mixture of *K* independent Poissons with mean (1 + *s*)*N/N*_*f*_ each, where *K* is the random number of ears chosen to replant, which is itself Poisson with mean *μ*_*K*_. By Poisson additivity, the numbers of local and migrant offspring are Poisson, with means *λ*_*P*_ = *λ*_*P*__*L*_ + *λ*_*P*__*M*_ and *μ*_*K*_ = *μ*_*KL*_ + *μ*_*KM*_ respectively. With probability *p*_*e*_, *λ*_*P*__*M*_ = *m*_*g*_(1 + *s*) and *μ*_*K*_ = *λ*_*P*__*L*_ = 0. Otherwise, with probability (1 − *p*_*e*_)(1 − *p*_*m*_), *μ*_*KL*_ = *N*_*f*_ */N* and *λ*_*P*__*L*_ = (1 + *s*)(1 − *m*_*g*_); and with probability (1 − *p*_*e*_)*p*_*m*_, *μ*_*KL*_ = (1 − *r*_*m*_)*N*_*f*_ */N* and *λ*_*P*__*L*_ = (1 − *r*_*m*_)(1 + *s*)(1 − *m*_*g*_). The migrant means are, with probability (1 − *p*_*e*_)*p*_*e*_*/*(1 − *p*_*e*_) = *p*_*e*_, *μ*_*KM*_ = *N*_*f*_ */N* and *λ*_*P*__*M*_ = 1 + *s*; while with probability (1 − *p*_*e*_)*p*_*m*_, *μ*_*KM*_ = *r*_*m*_*N*_*f*_ */N* and *λ*_*P*__*M*_ = (1 + *s*)(*r*_*m*_(1 − *m*_*g*_) + *m*_*g*_), and otherwise *μ*_*KM*_>_*>*_=0 and *λ*_*P*__*M*_= *m*_*g*_(1 + *s*).

**TABLE A1.** Parameter estimates used in calculations, and other notation.

|  |  |  |
| --- | --- | --- |
| complete seed stock replacement prob | $p_e$ | 0.068 |
| pollen migration rate | $m_g$ | 0.0083 |
| number of plants per field | $N$ | $10^5$ |
| number of ears used to replant | $N_f$ | 561 |
| mean ears per plant | $\mu_E$ | 3 |
| partial stock replacement prob | $p_m$ | 0.02 |
| mean proportion stock replaced | $r_m$ | 0.2 |
| pollen migration distance | $\sigma_p$ | 0 km |
| seed replacement distance | $\sigma_s$ | 50 km |
| distance between demes | $a$ | 15 km |
| width of altitudinal cline | $w$ | 62km |
| deleterious selection coefficient | $s_d$ | varies |
| beneficial selection coefficient | $s_b$ | varies |
| slope of selection gradient | $\alpha$ | $(s_d + s_b)/w$ |
| variance in offspring number | $\xi^2$ | varies |
| maize population density | $\rho$ | $5 \times 10^3$ |
| area of highland habitat | $A$ | 270,000 km<sup>2</sup> |
| mean dispersal distance | $\sigma$ | 1.8 km |

#### The generating function

The generating function of a Poisson with mean *λ* is *ϕ* (*u*; *λ*) = exp(*λ*(*u−*1)), and the generating function of a Poisson(*μ*) sum of Poisson(*λ*) values is *ϕ*(*ϕ* (*u*; *λ*); *μ*). Therefore, the generating function for the diploid process, ignoring spatial structure, is

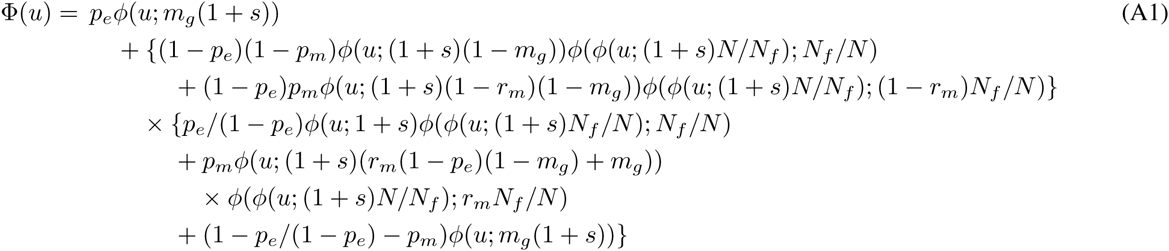

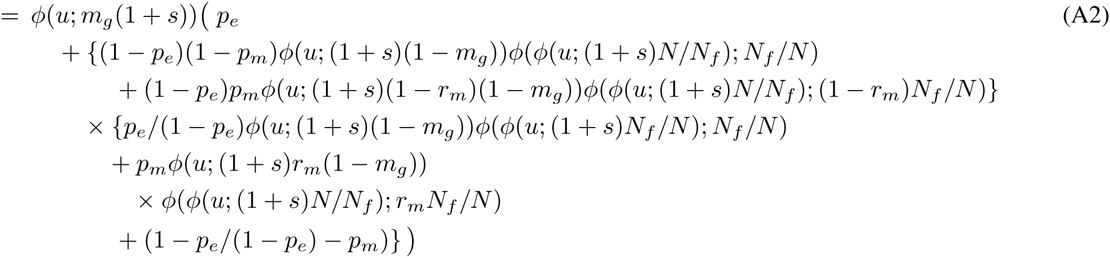

To get the generating function for a haploid, replace every instance of 1 + *s* by (1 + *s*)*/*2.

As a quick check, the mean total number of offspring of a diploid is

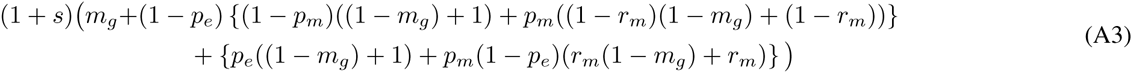

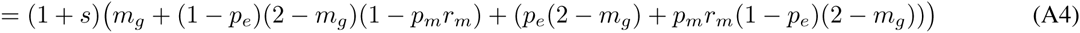

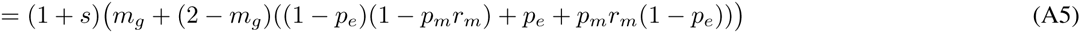

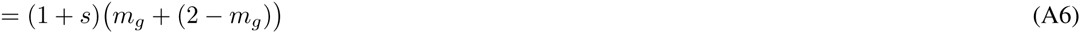

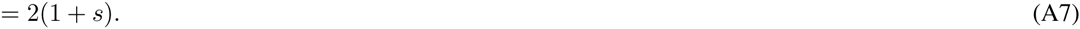

We show numerically later that the probability of establishment is very close to 2*s* over the variance in reproductive number (as expected). It is possible to write down an expression for the variance, but the exact expression does not aid the intuition.

#### Migration and spatial structure

To incorporate spatial structure, suppose that the locations *ℓ*_*k*_ are arranged in a regular grid, so that *ℓ*_*k*_ = *ak*. Recall that *s*_*k*_ is the selection coefficient at location *k*. If the total number of offspring produced by an individual at *ℓ*_*i*_ is Poisson(*λ*_*i*_), with each offspring independently migrating to location *j* with probability *m*_*ij*_, then the number of offspring at *j* is Poisson(*m*_*ij*_*λ*_*i*_), and so the generating function is

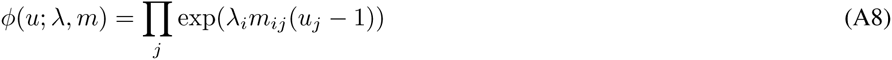

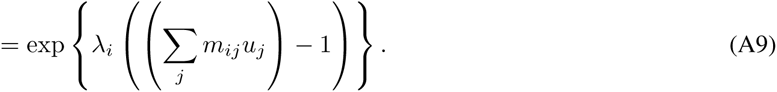

We can then substitute this expression into equation (A1), with appropriate migration kernels for pollen and seed dispersal.

For migration, we need migration rates and migration distances for both wind-blown pollen and for farmer seed exchange. The rates are parameterized as above; we need the typical dispersal distances, however. One option is to say that the typical distance between villages is *d*_*v*_, and that villages are discrete demes, so that pollen stays within the deme (pollen migration distance 0) and seed is exchanged with others from nearby villages; on average *σ*_*s*_ distance away in a random direction. The number of villages away the seed comes from could be geometric (including the possibility of coming from the same village).

#### Dispersal distance

The dispersal distance – the mean distance between parent and offspring – is equal to the chance of inter-village movement multiplied by the mean distance moved. This is

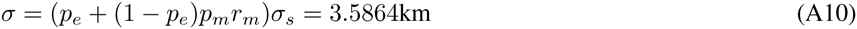

at the parameter values above.

Iterating the generating function above finds the probability of establishment as a function of distance along the cline. This is shown in figure A1. Note that the approximation 2*s* divided by the variance in offspring number is quite close.

In the main text, we used a rough upper bound on the rate of migration that ignored correlations in migrants. As we show in Ralph and Coop (2014a), the rate of adaptation by diffusive migration is more precisely

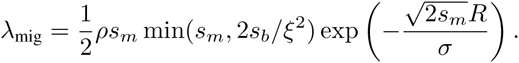

First note that for 10^−1^ *≤s*_*m*_ *≤*10^−4^, the value 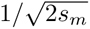 is between 2 and 70 – so the exponential decay of the chance of migration falls off on a scale of between 2 and 70 times the dispersal distance. Above we have estimated the dispersal distance to be *σ ≈* 3.5 km, and far below the mean distance *σ*_*s*_ to the field that a f<u>arm</u>er replants seed from, when this happens, which we have as *σ*_*s*_ = 50 km. Taking *σ* = 3.5 km, we have that 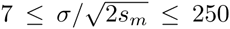 A very conservative upper bound might be *σ ≤ σ*_*s*_/10 (if farmers replaced 10% of their seed with long-distance seed every year). At this upper bound, we would have 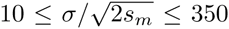, which is not very different. This makes the exponential term small since *R* is on the order of thousands of kilometers.

Taking *σ* = 3.5 km, we then compute that if *s*_*m*_ = 10^−4^ (very weak selection in the lowlands), then for *R* = 1, 000 km, the migration rate is *λ*_mig_ ≤10^−5^, i.e. it would take on the order of 100,000 generations (years) to get a successful migrant only 1,000 km away, under this model of undirected, diffusive dispersal. For larger *s*_*m*_, the migration rate is much smaller.

#### Migration rate of deleterious alleles

In the main text we computed *λ*_mig_, the rate at which new adaptive alleles appeared by mutation. A corresponding expression for the chance that an allele moves from one highland population to another is harder to intuit. This problem is studied in more depth in Ralph and Coop (2014a), under the assumption that the alleles are deleterious between the highlands. Since such deleterious alleles are much less likely to transit than neutral ones, the analysis in the main text implies that gene flow is unlikely to have shared these alleles between highland regions. However, because spatially continuous models assuming selective effects are better understood than neutral ones, and we do expect a tradeoff between highland- and lowland-adaptation, it is useful to understand what happens in this case as well.

If an allele is beneficial at high elevation and fixed in the Mesoamerican highlands but is deleterious at low elevations, then at equilibrium it will be present at low frequency at migration-selection balance in nearby lowland populations (Haldane 1948; Slatkin 1973). This equilibrium frequency decays exponentially with distance, so that the highland allele is present at distance *R* from the highlands at frequency *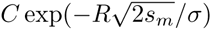* where *s*_*m*_ is the deleterious selection coefficient for the allele in low elevation, *σ* is the mean dispersal distance, and *C* is a constant depending on geography (*C ≈* 1*/*2 is close). Multiplying this frequency by a population size gets the predicted number (average density across a large number of generations) of individuals carrying the allele. Therefore, in a lowland population of size *N* at distance *R* from the highlands, 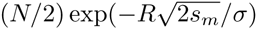 is equal to the probability that there are any highland alleles present, multiplied by the expected number of these given that some are present. Since we assume the allele is deleterious in the lowlands, if *R* is large there are likely none present; but if there are, the expected number is of order 1*/s*_*m*_ (Geiger 1999; Ralph and Coop 2014a). This therefore puts an upper bound on the rate of migration of

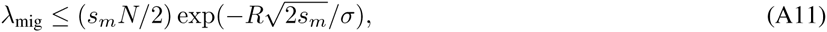

and we we would need to wait *T*_mig_ = 1*/λ*_mig_ generations for a rare such excursion to occur. This calculation omits the probability that such an allele fixes (≈2*s*_*b*_*/ξ*^2^) (discussed above) and the time to reach migration-selection balance (discussed in the main text); both of these omissions mean we underestimate *T*_mig_.

##### Results for gene flow of deleterious alleles

From our demographic model we have estimated a mean dispersal distance of *σ* ≈3.5 kilometers per generation. With selection against the highland allele in low elevations 10^−1^ *≥ s*_*m*_ *≥* 10^−4^, the distance *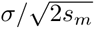* over which the frequency of a highland-adaptive, lowland-deleterious allele decays into the lowlands is still short: between 7 and 250 kilometers. Since the Mesoamerican and Andean highlands are around 4,000 km apart, the time needed for a rare allele with weak selective cost *s*_*m*_ = 10^−4^ in the lowlands to transit between the two highland regions is *T*_mig_ ≈ 8 × 10^4^ generations. While the exponential dependence on distance in equation (A11) means that shorter distances could be transited more quickly, the waiting time *T*_mig_ is also strongly dependent on the magnitude of the deleterious selection coefficient: with *s*_*m*_ = 10^−4^, *T*_mig_ ≈ 25 generations over a distance of 2,000 km, but increases to ≈ 10^8^ generations with a still weak selective cost of *s*_*m*_ = 10^−3^.

---

**Table S1.**
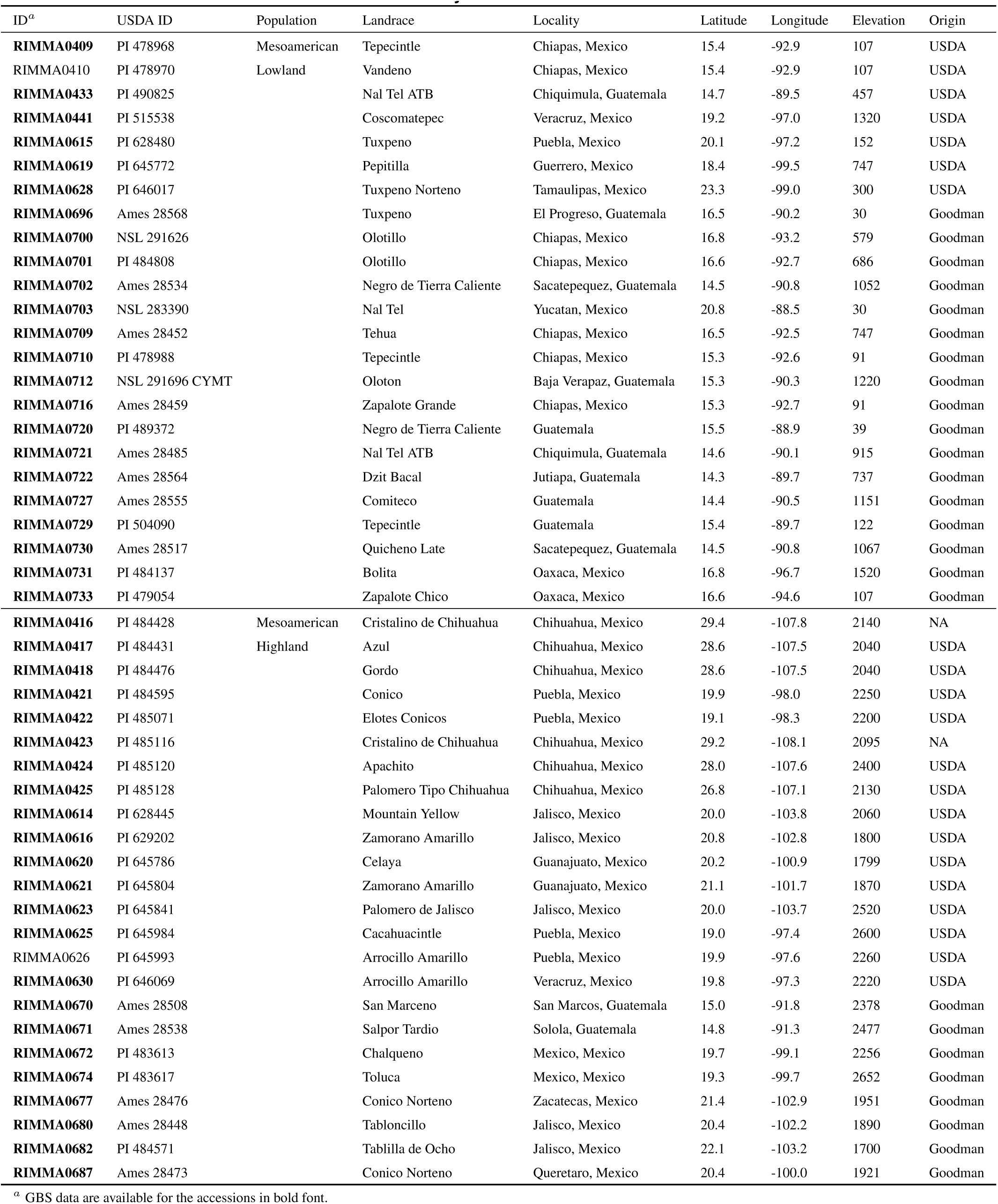
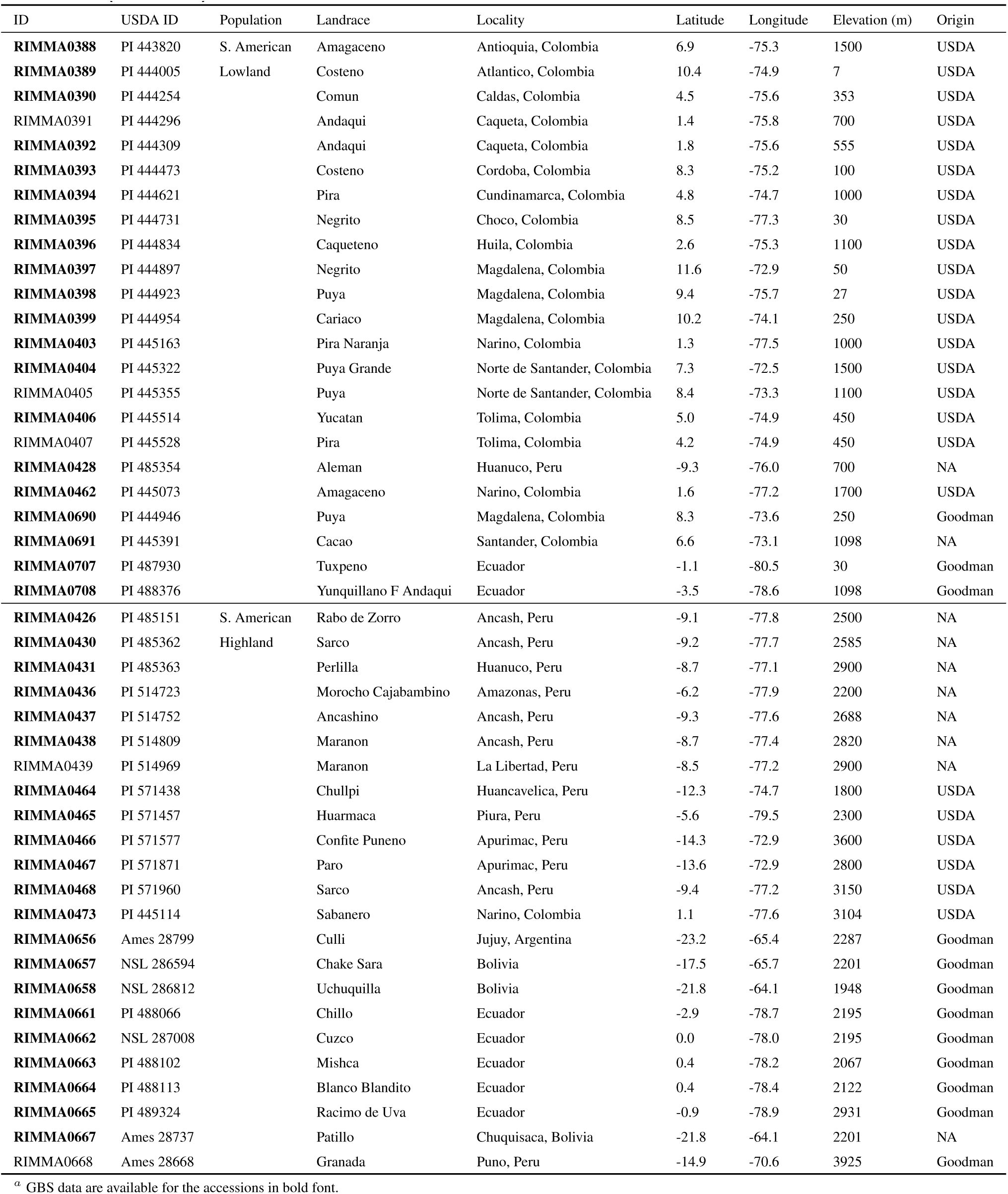
List of maize landraces used in this study.

**Table S2.** Patterns of adaptation.

| Population | Pattern of adaptation | No. of SNPs | No. of SNPs supported by PHS test | Significance <sup>a</sup> |
| --- | --- | --- | --- | --- |
| Mesoamerica | Highland adaptation | 264 | 172 (65.2%) | $P < 10^{-3}$ |
| | Lowland adaptation | 101 | 66 (65.3%) | $P < 0.05$ |
| S. America | Highland adaptation | 164 | 230 (71.3%) | $P < 10^{-5}$ |
| | Lowland adaptation | 70 | 50 (71.4%) | $P < 0.05$ |

<sup>a</sup> Probability of the observed percent of SNPs showing a lower empirical quantile. Under neutrality, 50% of SNPs should have lower PHS values in the focal population; higher values indicate evidence of selection. See the main text for details.

**Table S3.** List of metabolic pathways showing evidence of convergent adaptation

|  |
| --- |
| Colanic acid building blocks biosynthesis |
| Purine nucleotides *de novo* biosynthesis II |
| Adenosine nucleotides *de novo* biosynthesis |
| NAD/NADH phosphorylation and dephosphorylation |
| tRNA charging pathway |
| Superpathway of phenylalanine biosynthesis |
| Superpathway of tryptophan biosynthesis |
| Aspartate biosynthesis |
| Tryptophan biosynthesis |
| Glutamine biosynthesis III |
| Isoleucine biosynthesis I |
| Threonine biosynthesis |
| Galactose degradation III |
| UDP-glucose biosynthesis (from glucose 6-phosphate) |
| Triacylglycerol biosynthesis |
| Phospholipid biosynthesis II |
| Phosphatidylglycerol biosynthesis I (plastidic) |
| Phosphatidylglycerol biosynthesis II (non-plastidic) |
| CDP-diacylglycerol biosynthesis II |
| CDP-diacylglycerol biosynthesis I |
| Ethylene biosynthesis from methionine |
| Stachyose degradation |
| Homogalacturonan degradation |
| Betanidin degradation |
| Aspartate degradation II |
| Phosphate utilization in cell wall regeneration |
| Phosphate acquisition |
| Superpathway of cytosolic glycolysis (plants), pyruvate dehydrogenase and TCA cycle |
| C4 photosynthetic carbon assimilation cycle |
| Glycolysis IV (plant cytosol) |
| Glycolysis I |
| Glycolysis III |

**Table S4.**
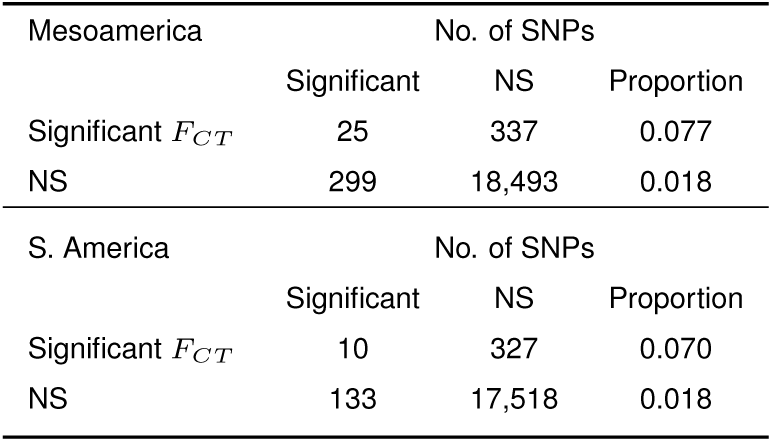
*F*_*CT*_ between *parviglumis* and *mexicana*.

**Table S5.** *F*_*ST*_ outlier SNPs and *mexicana* introgression.

| Introgression status | Population | $F_{ST}$ outlier SNPs | All other SNPs |
| --- | --- | --- | --- |
| Introgressed | Mesoamerica | 114 | 1953 |
|  | S. America | 26 | 1721 |
| Not introgressed | Mesoamerica | 558 | 73892 |
|  | S. America | 379 | 60666 |

**Figure S1.**
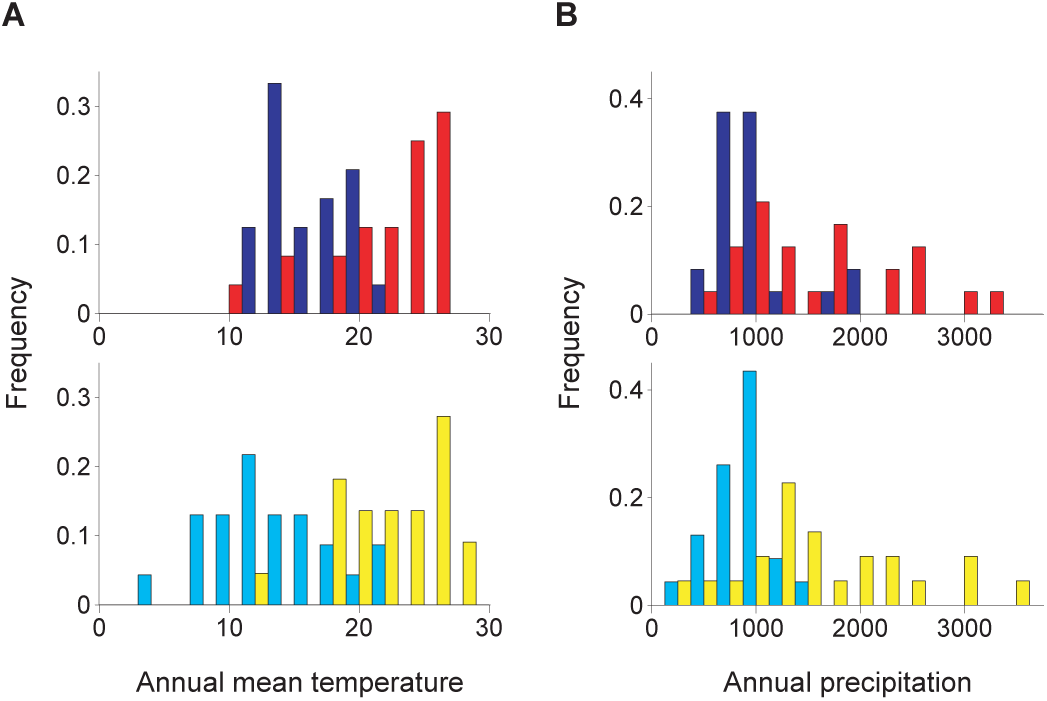
Annual mean temperature and annual precipitation of the locations of the maize samples used in this study. Red, blue, yellow and light blue bars represent Mesoamerican lowland, Mesoamerican highland, S. American lowland and S. American highland populations, respectively.

**FIGURE S2.**
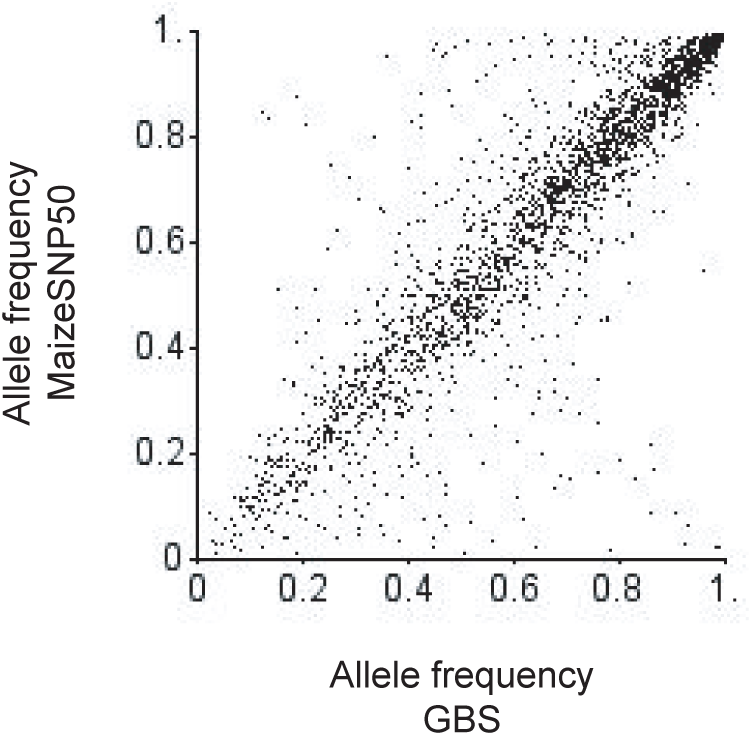
Correlation of allele frequencies between GBS and MaizeSNP50 data. We used overlapping SNPs with *n ≥* 40 for both data sets. The correlation coefficient is 0.890 (*P* < 10^−5^ by permutation test with 10^5^ replications).

**FIGURE S3.**
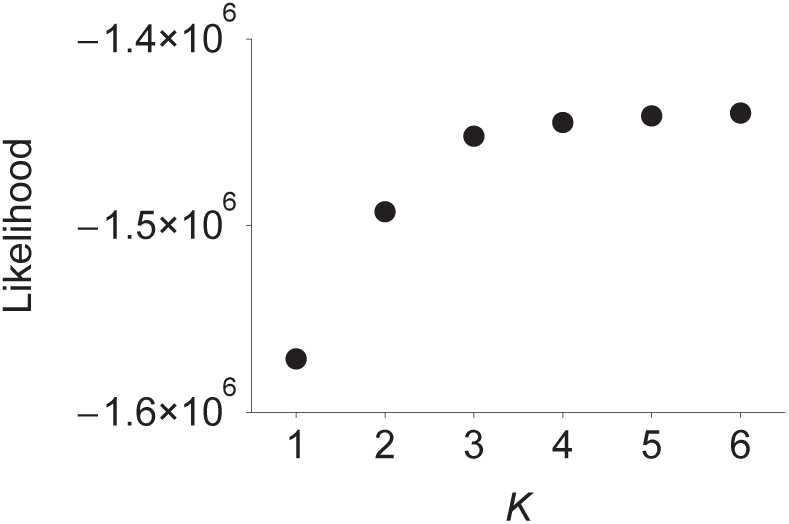
Likelihood of STRUCTURE analyses given the number of populations *K*.

**FIGURE S4.**
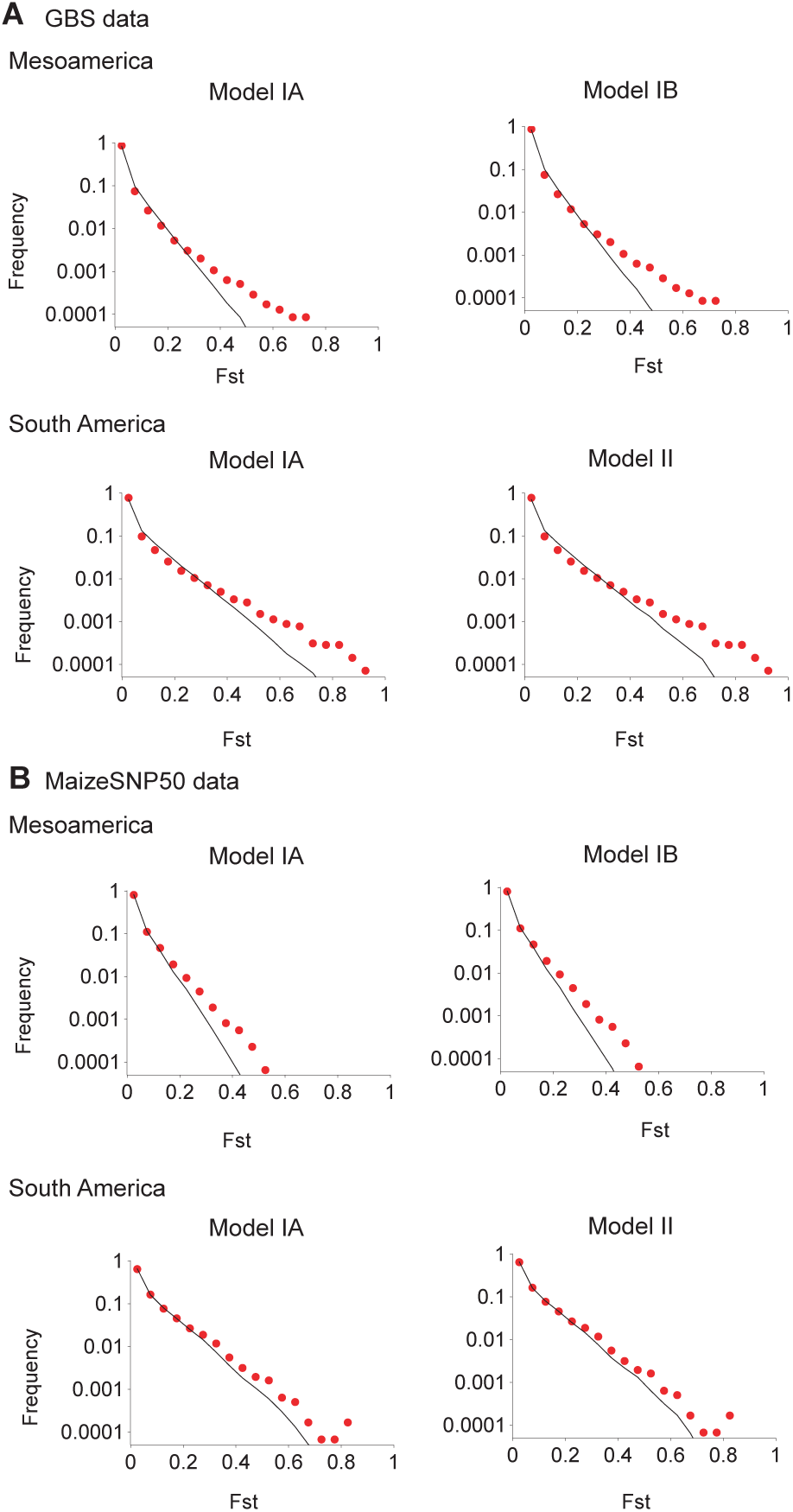
Observed and expected distributions of *F*_*ST*_ values in GBS (A) and MaizeSNP50 data (B). The *y*-axes represent the expected (solid lines) and observed (red dots) frequency of SNPs for a range of *F*_*ST*_ values in bins of 0.05.

**FIGURE S5.**
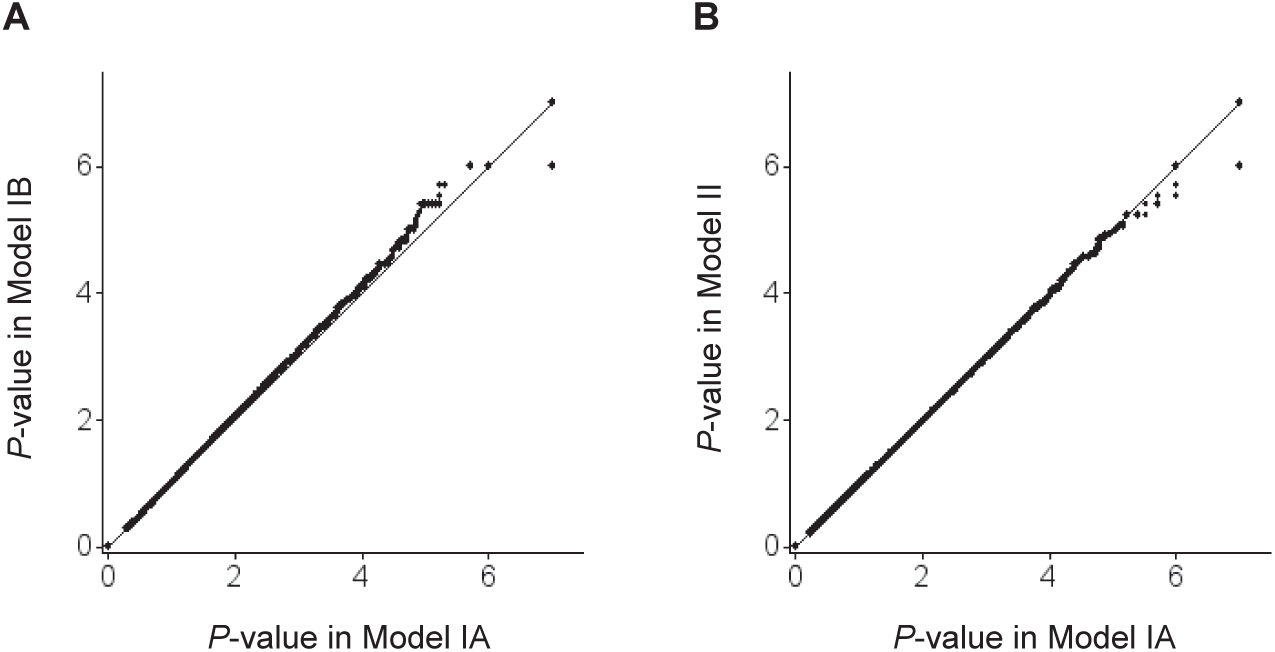
Q-Q plot for log_10_-scaled *P*-values of population differentiation between lowland and highland populations. (A) Model IA *v.s.* Model IB in Mesoamerica, (B) Model IA *v.s.* Model II in S. America.

**FIGURE S6.**
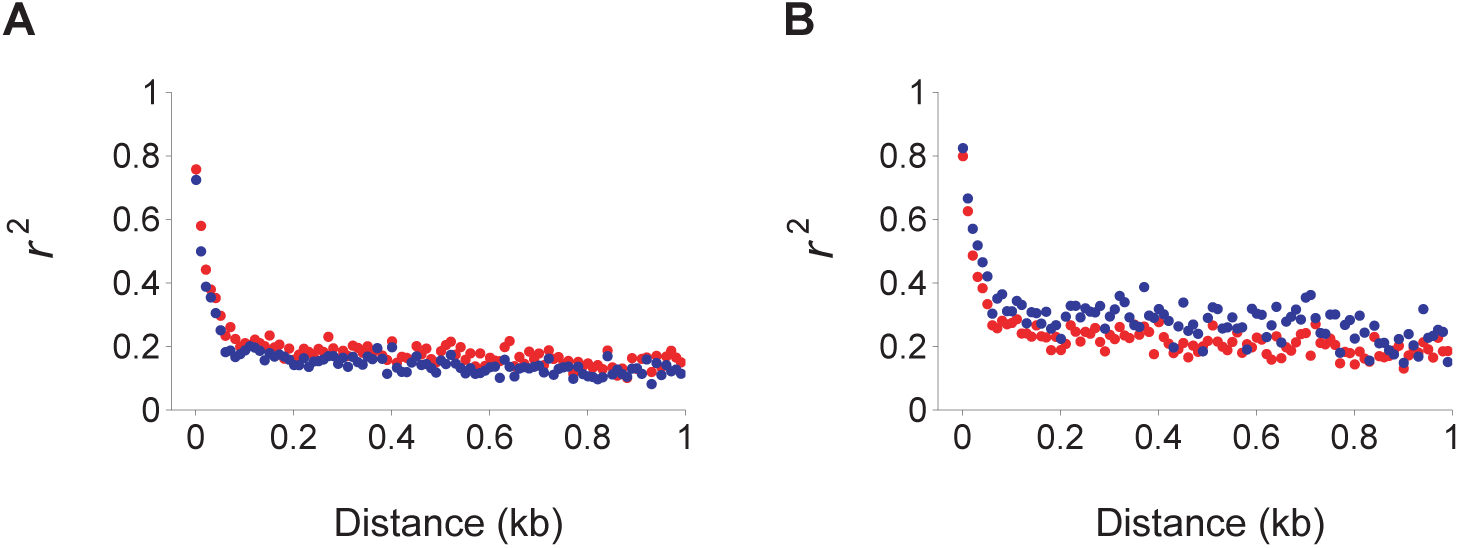
Pattern of decay of linkage disequilibrium in Mesoamerica (A) and S. America (B). Red and blue dots represent lowland and highland populations, respectively. *r*^2^ values are reported as averages within 10-bp distance bins.

